# Targeted degradation of CTCF decouples local insulation of chromosome domains from higher-order genomic compartmentalization

**DOI:** 10.1101/095802

**Authors:** Elphège P. Nora, Anton Goloborodko, Anne-Laure Valton, Johan Gibcus, Alec Uebersohn, Nezar Abdennur, Job Dekker, Leonid A. Mirny, Benoit G. Bruneau

## Abstract

The molecular mechanisms underlying folding of mammalian chromosomes remain poorly understood. The transcription factor CTCF is a candidate regulator of chromosomal structure. Using the auxin-inducible degron system in mouse embryonic stem cells, we show that CTCF is absolutely and dose-dependently required for looping between CTCF target sites and segmental organization into topologically associating domains (TADs). Restoring CTCF reinstates proper architecture on altered chromosomes, indicating a powerful instructive function for CTCF in chromatin folding, and CTCF remains essential for TAD organization in non-dividing cells. Surprisingly, active and inactive genome compartments remain properly segregated upon CTCF depletion, revealing that compartmentalization of mammalian chromosomes emerges independently of proper insulation of TADs. Further, our data supports that CTCF mediates transcriptional insulator function through enhancer-blocking but not direct chromatin barrier activity. These results define the functions of CTCF in chromosome folding, and provide new fundamental insights into the rules governing mammalian genome organization.

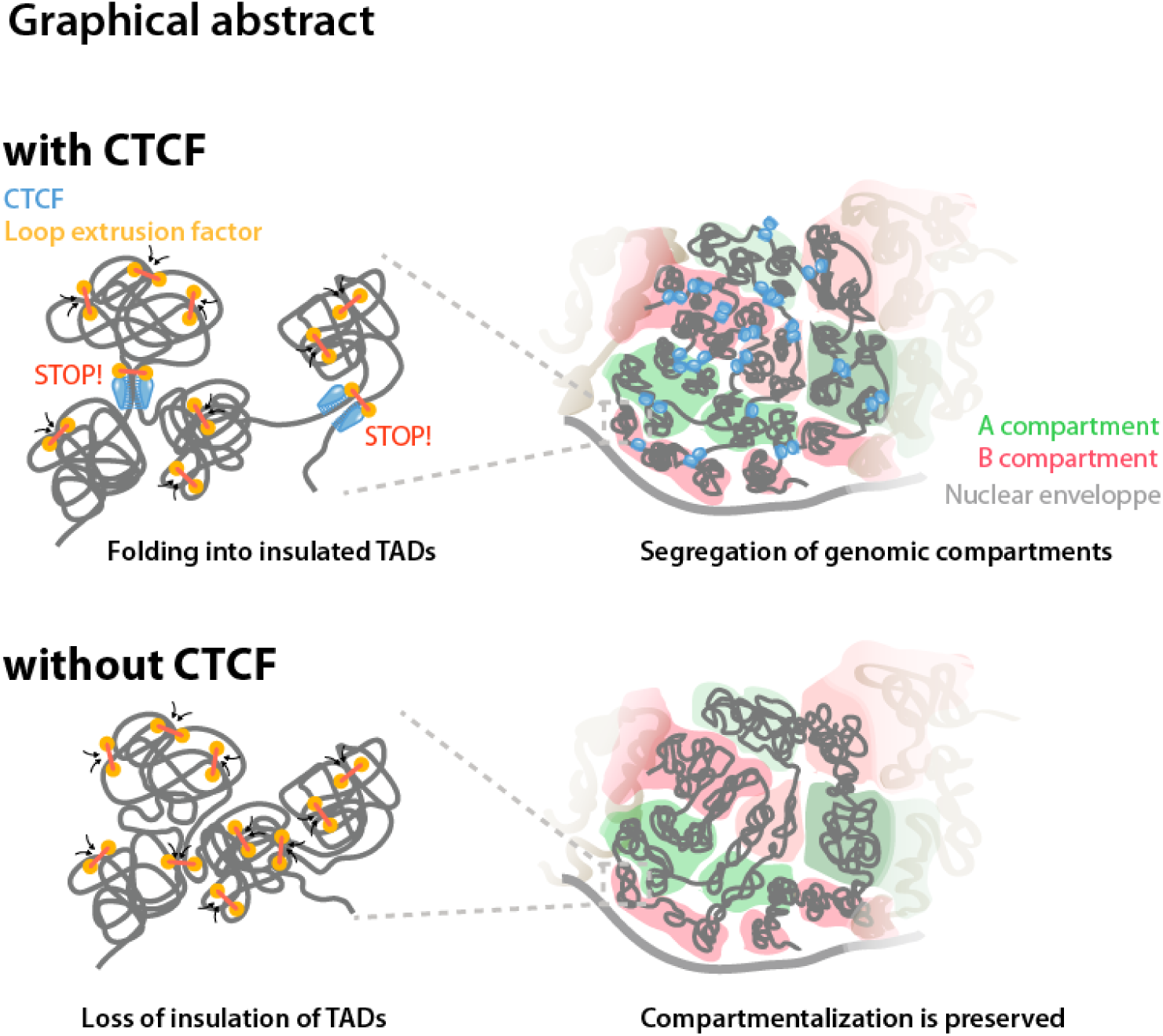

## INTRODUCTION

Chromosomes meet the dual challenge of packaging DNA into the nucleus and, at the same time, enabling access to genetic information. Decades of work on chromosome organization have tackled the link between chromosome structure and genetic functions (Belmont, 2014; Cremer et al., 2015). Patterns of genome folding have been scrutinized with ever-increasing precision, but the identity and roles of the underlying molecular actors are still poorly understood, limiting our functional understanding of chromosome architecture. While genome organization and molecular actors differ between distant species (Cubeñas-Potts et al., 2016; Dekker and Heard, 2015; Ea et al., 2015), here we focus on mammals.

Mammalian chromosomes are profoundly heterogeneous. Euchromatin comprises open chromatin fibers and gene-rich regions (Gilbert et al., 2004) while heterochromatin is condensed, gene-poor and transcriptionally dormant. This highlights the remarkable correlation between the cytological, biochemical and sequence organization of chromosomes. Chromosomes can be further segmented into domains belonging to two main types of spatial compartments, as reveled by high-throughput Chromosome Conformation Capture (3C), with chromatin contacts being more frequent between loci of the same compartment type, both within and between chromosomes (Lieberman-Aiden et al., 2009). When reported on linear genomic maps, the alternating pattern of compartment types forms a domain-wide arrangement that aligns strikingly with regional chromatin states (Bickmore and van Steensel, 2013; Bonev and Cavalli, 2016). The euchromatic A- compartment contains most actively transcribed regions, while the B-compartment corresponds to megabase-sized gene-poor Lamina-Associated Domains (LADs (Guelen et al., 2008; Kind et al., 2015)) which replicate late in S-phase (Hiratani et al., 2008; Ryba et al., 2010).

At a more local scale, chromosomes are partitioned into sub-megabase segments that tend to self-associate and thus are relatively insulated from neighboring domains, forming structures termed Topologically Associating Domains (TADs, (Dixon et al., 2012; Nora et al., 2012)). The borders of TADs are frequently demarcated by the binding of the CCCTC-binding factor (CTCF), (Dixon et al., 2012; Phillips-Cremins et al., 2013) a broadly expressed zinc-finger nucleic acid binding protein initially that is involved in transcriptional insulation and chromosomal interactions (Ghirlando and Felsenfeld, 2016; Merkenschlager and Nora, 2016). Ultra-high resolution Hi-C analyses demonstrated the existence of a peak of 3C signal between some CTCF-bound boundaries of TADs, referred to as contact domains at this scale – indicative of interaction through chromatin looping (Rao et al., 2014). Deleting a TAD boundary, or even just the underlying CTCF site, can lead to loss of physical insulation and subsequent encapsulation of the two abutting TADs in a single domain (Lupiáñez et al.; Narendra et al., 2015; Nora et al., 2012; Sanborn et al., 2015; Tsujimura et al., 2015). This highlights the crucial role of boundaries in mediating the physical insulation of neighboring chromosome domains, with important implications for disease-causing chromosomal rearrangements in humans (Flavahan et al., 2016; Franke et al., 2016; Hnisz et al., 2016).

Strikingly, in most of the cases, a pair of CTCF sites only engage in contact above local background if they are in a convergent linear orientation (Rao et al., 2014), creating an asymmetry in the insulation pattern (Vietri Rudan et al., 2015). This arrangement is important: inverting a single CTCF site can be enough to rewire the direction of looping and disrupt proper packaging of the underlying chromosomal segment into an insulated TAD (Guo et al., 2015; Lupiáñez et al.; Sanborn et al., 2015; de Wit et al.). Polymer modelling studies have proposed that CTCF mediates TAD insulation by acting as a polar blocking factor to Cohesin translocation along the DNA during the formation and expansion of chromatin loops (Fudenberg et al., 2016; Sanborn et al., 2015).

Locus-specific studies have implicated the CTCF protein itself in mediating chromosome folding (Splinter, 2006). Yet, genome-wide assays after RNAi revealed only very limited consequences of CTCF depletion – although qualitatively concordant with site deletion experiments (Zuin et al., 2014). Genetic manipulation of CTCF has proven difficult as it is essential for development (Moore et al., 2012; Sleutels et al., 2012; Soshnikova et al., 2010; Wan et al., 2008) and proliferation of cultured cells (González-Buendía et al., 2014), hampering the understanding of the exact role of CTCF in mammalian chromosome folding and genome functions. It is currently unclear to what extent CTCF is actually required for chromatin architecture and which levels of genome organization this factor controls.

Here we used a conditional degradation strategy in mouse embryonic stem cells (mESCs), the Auxin-Inducible Degron (AID) system (Nishimura et al., 2009), to acutely and reversibly deplete CTCF below detectable levels. We demonstrate that CTCF is a major determinant of mammalian chromosome folding. Its role is however restricted to sub-megabase genome organization, with loss of CTCF leading dose-dependently to insulation defects at TAD boundaries and abrogating long-range contact between CTCF sites. Importantly, CTCF depletion did not disrupt A/B compartments, revealing that local insulation and higher-order compartmentalization must rely on distinct molecular determinants. Our observations also reveal an important activator effect of CTCF through direct promoter binding, support a role for CTCF as an enhancer blocker, and refute its proposed function as a direct barrier to H3K27me3 spreading.

## RESULTS

### Acute CTCF Depletion with the Auxin-Inducible Degron System

To deplete endogenous CTCF in mESCs, we leveraged the AID system that enables rapid and acute protein degradation in a reversible manner (Nishimura et al., 2009). We targeted the stop codon of both *Ctcf* alleles to introduce a 44-amino acid version of the AID tag (residues 71–114, (Morawska and Ulrich, 2013)) with an eGFP cassette, in male mESCs (Figure 1A, Supplementary table1). We also introduced a transgene encoding the Tir1 F-box protein from *Oryza sativa* (rice), which can bind to the AID peptide only in the presence of auxin, triggering polyubiquitination and proteasome-dependent degradation (Tan et al., 2007). The resulting cell line is referred to as CTCF-AID hereafter.

**Figure 1.**
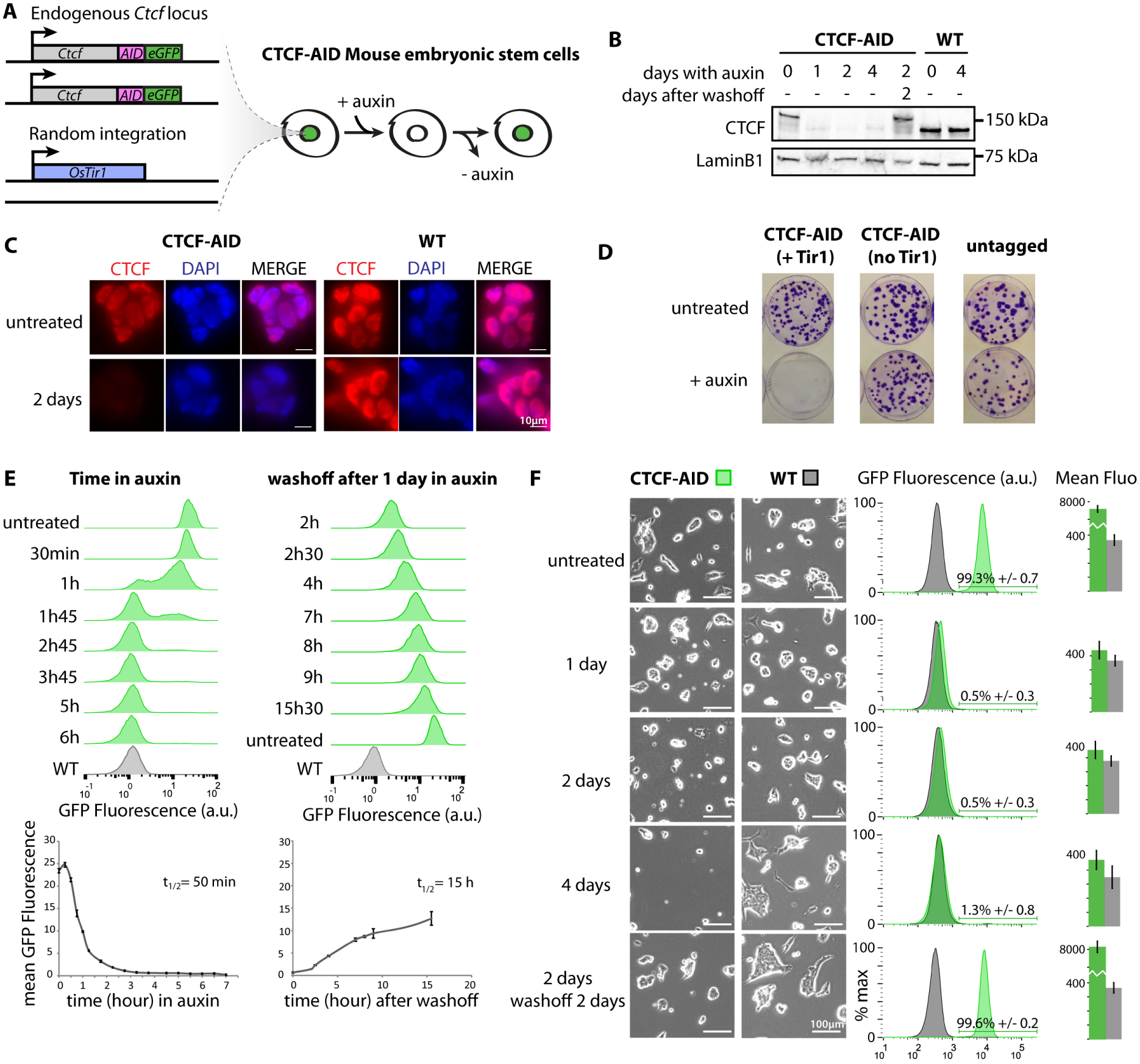
Acute and reversible depletion of CTCF with the auxin-inducible degron system in mESCs. (A)Deployment of the AID system at *Ctcf* in mESCs. (B) Western-blot analysis showing reversible loss of CTCF in the CTCF-AID cells. (C) Immunofluorescence analysis of CTCF-AID mESCs. (D) Survival is only compromised in CTCF-AID cells treated with auxin after introduction of the OsTir1 F-box transgene. (E) Time-course flow-cytometry after addition of auxin or washoff after 1 day of treatment. (F) Brightfield images of mESC colonies after auxin treatment demonstrating cells tolerate a 2-day depletion with no adverse effects on viability. **See also Figure S1**

Adding auxin to the culture medium (see methods) depleted CTCF to levels that could not be detected by Western blot, and washing out auxin allowed CTCF to accumulate back to initial levels (**Figure 1B**). Auxin in itself was neutral to untagged mESCs (**Figure 1D and E**), and RNA-seq revealed no differential gene activity after treating untagged cells for up to 4 days (**Supplementary tables 2 and 3**). As reported previously the AID fusion led to slight constitutive destabilization (Morawska and Ulrich, 2013), so that basal CTCF levels were about 2–3-fold less in the AID-eGFP fusion line compared to the untagged parental line (**Figure 1B and C**). RNA-seq revealed 72 differentially expressed genes between the parental and untreated CTCF-AID lines (**Supplementary table 3**). Cells could nevertheless be expanded and subcloned normally (**Figure 1D**), indicating that the AID-eGFP fusion does not abrogate the essential functions of CTCF. In contrast, auxin-mediated degradation of CTCF prevented subcloning of CTCF-AID cells, recapitulating the full CTCF knockout phenotype in mESCs (Sleutels et al., 2012) (**Figure 1D**).

Time-course flow-cytometry demonstrated maximal depletion of CTCF-AID as early as 3:45 hours after adding auxin (**Figure 1E**). Recovery initiated readily after washoff and was half complete by 15h (**Figure 1E**). Interestingly, acute CTCF depletion in mESCs was tolerated for 2 days without obvious cell death or differentiation (**Figure 1F**). Depleting for longer than 2 days slowed cell proliferation darmatically (**Figure 1F and S1A-B**). Importantly, CTCF depletion in mESCs did not block cells in a specific phase of the cell cycle, and did not induce DNA damage or aneuploidy (**Supplementary figures 1 D-E**). Cell death increased after 4 days of depletion but we did not observe massive apoptosis (**Figure S1**), contrasting with other cellular contexts (Soshnikova et al., 2010; Watson et al., 2014). Our system can therefore be used during at least two days after auxin addition (3-4 cell divisions) to study the immediate consequences of acute CTCF depletion without adverse effect on cell survival and proliferation.

### Auxin Treatment Severely Depletes CTCF from Chromatin

CTCF binding patterns, as measured by ChIP-exo in untreated CTCF-AID mESCs, were highly similar to untreated or 2-day treated WT untagged cells, highlighting that auxin treatment in itself does not affect CTCF binding, nor does tagging with the AID-eGFP cassette (**Figure S2A**). Using ChIP-seq in CTCF-AID cells after 2 days of auxin we detected only 27% of the initial CTCF peaks. (**Figure S1B-C and supplementary table 4**). The enrichment level in persistent peaks was severely reduced (**Figure S2D-E**), indicating that CTCF occupancy is lost or considerably lower at all of its genomic target sites after depletion. Persistent peaks correspond to the sites of highest occupancy in untreated cells, which may arise from the few cells that do not respond to auxin (0.5–1% of the population, **Figure 1F**) or from binding to the highest affinity sites by the very small amount of residual CTCF. ChIP-seq patterns from cells where auxin was washed off after a 2-day treatment was virtually identical to untreated cells, revealing that CTCF readily regains access to all of its cognate binding sites after transient depletion in mESCs (**Figure S2B-E**). Finally, depletion efficiency was equally efficient irrespective of local binding site density, suggesting that large-scale genomic clustering of binding sites does not promote cooperative binding for CTCF (**Figure S2F-H**).

**Figure 2.**
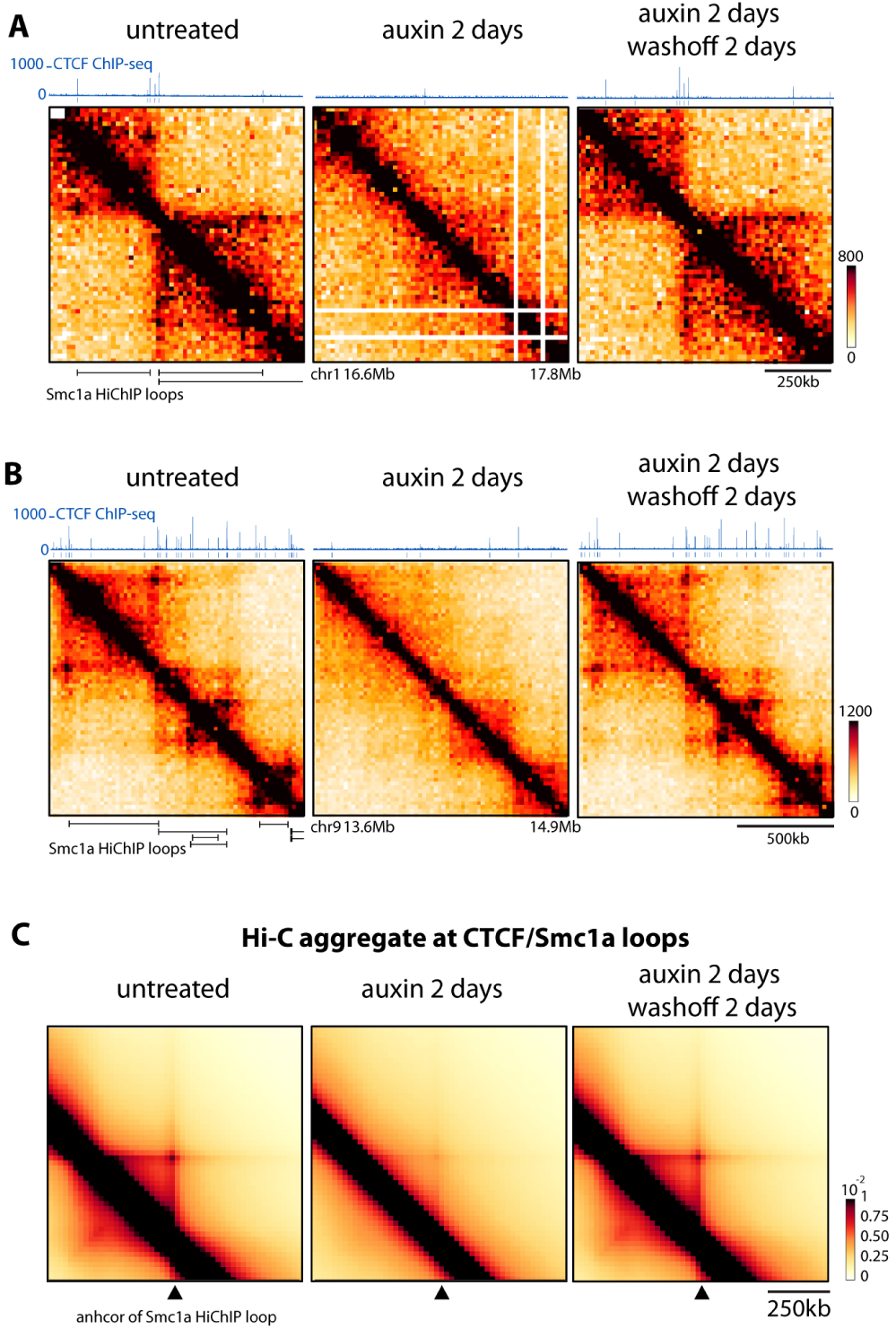
CTCF is required for accumulating loops between CTCF/Cohesin binding sites. (A-B) Loss of Hi-C peaks upon CTCF depletion. 1.5 Mb snapshot of Hi-C data at 20kb resolution CTCF-AID mESCs aligned with CTCF ChIP-seq and the Smc1a HiChIP loops identified by Mumbach et al. 2016. Normalized Hi-C counts are multiplied by 10^5^. (C) Genome-wide aggregation of normalized Hi-C signal anchored at Smc1a HiChip loops separated by 280 to 380kb (1196 loops). Similar results were obtained for smaller and larger loops. **See also Figure S2**

### CTCF is required for formation of chromatin loops at CTCF sites

In order to measure changes in chromosome organization upon CTCF depletion we performed high-throughput 3C-based experiments. Current technologies require extremely deep sequencing to interrogate changes in contact frequencies between individual genomic loci below the megabase scale at the genome-wide scale. Therefore, we first focused on the the *X-inactivation centre* locus (*Xic*) using 3C Carbon-Copy (5C, (Dostie et al., 2006)), with our male undifferentiated mESCs (which harbor a single active X chromosome). The *Xic* displays strong well-characterized CTCF-anchored interactions between the *Linx, Chicl* and *Xite* elements (Giorgetti et al., 2014; Nora et al., 2012) readily detected by 5C (**Figure S2I and Supplementary Table 5**). Chromosomal organization at the *Xic* in the untagged parental line was not perturbed by auxin and was identical in the untreated CTCF-AID (**Figure S2J**). In contrast, auxin-mediated depletion of CTCF led to the complete disappearance of these 5C peaks, demonstrating they depend on CTCF (**Figure S2I**). Auxin washoff reset these chromosomal contacts to a pattern that is indistinguishable from the initial untreated sample.

To extend these observations to the entire genome, we performed Hi-C in untreated, 2-day treated cells as well as after a 2-day washoff. Our 20kb resolution data (**Figure 2A-B**) did not allow us to perform robust *de novo* calling of loops. However, given that most CTCF binding events overlap with Cohesin enrichment by ChIP-seq (Parelho et al., 2008; Rubio et al., 2008; Wendt et al., 2008), we performed a meta-analysis of loops by aggregating our Hi-C signal at CTCF/Cohesin bound loops, as previously detected by high-resolution HiChip for Smc1a in mESCs (Mumbach et al., 2016). This confirmed that CTCF is required for the interaction between CTCF/Cohesin bound loop-anchor loci genome-wide, and that bringing CTCF back is sufficient to restore these specific contacts (**Figure 2C**).

### CTCF Depletion Triggers Dramatic Loss of TAD Insulation

We next investigated the integrity of TAD folding upon CTCF depletion. Our Hi-C maps revealed extensive ectopic contacts across initial TAD boundaries, clearly visible as early as 24 h after CTCF depletion by 5C at the *Xic* (**Figure 3A**). These changes were again fully reversible: washing off the auxin for 2 days after a 2-day treatment led to refolding of chromatin into properly insulated TADs (**Figure 3**).

**Figure 3.**
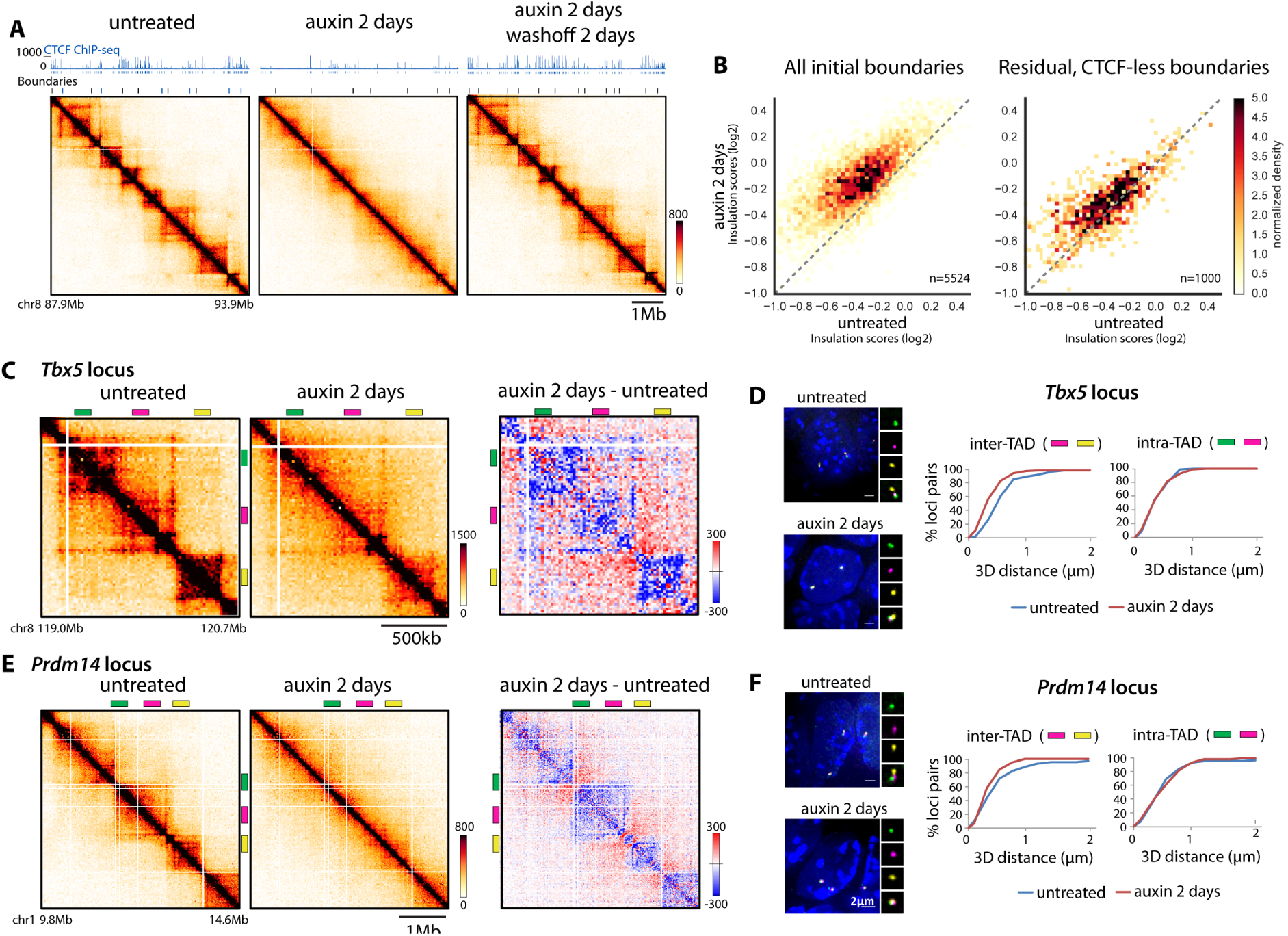
CTCF instructs insulation of TADs. (A) A 6Mb snapshot of Hi-C data at 20kb resolution from CTCF-AID mESCs aligned with CTCF ChIP-seq highlighting disrupted segmental folding into TADs upon CTCF depletion. Normalized Hi-C counts are multiplied by 10^5^. (B) Left: CTCF depletion dampens insulation at TAD boundaries (higher insulation score over 100kb surrounding boundaries). Right: residual boundaries detected after CTCF depletion (and without persistent CTCF peaks, ^~^20% of total boundaries) maintain insulation independently of CTCF. Note that lower score denotes higher insulation potential. (C) Snapshot of Hi-C data at the *Tbx5* locus and differential contact map showing more inter-TAD (red) and fewer intra-TAD (blue) contacts are detected after CTCF depletion. (D) 3D distance measurement from DNA FISH highlighting that CTCF depletion triggers inter-TAD compaction but does not affect intra-TAD packaging at the cytological level (E-F) same as C-D at the *Prdm14* locus. **See also Figure S3**

To quantify this behavior genome-wide and identify loci that may deviate from it, we scored insulation potential across all chromosomes using our Hi-C data (Crane et al., 2015)(**Supplementary Table 6 and Methods**). Our resolution enabled calling 5524 boundaries for a median TAD size of 340 kb (mean of 450kb) in untreated cells. Loss of CTCF led to loss of insulation at most boundaries (>80%, **Figure 3B**). This highlights that CTCF is not only required for local looping between TAD boundaries (**Figure 2**) but also for preventing the chromosomal segments enclosed by these loops from engaging with their neighboring genomic regions across the TAD boundary (**Figure 3**).

A subset of boundaries persisted after CTCF depletion. After removing those that displayed residual CTCF binding by ChIP-seq we identified 1000 persistent CTCF-less boundaries (18% of initial boundaries). Insulation of these persistent boundaries was much lss affected by CTCF depletion (**Figure 3B**).

To explore how changes measured by Hi-C translate at the cytological level, we used 3D DNA Fluorescent *in situ* Hybridization (FISH) to simultaneously measure the nuclear distance between three probes. Two probes were in the same TAD, and the third was separated by one or more TAD boundaries that lose insulation potential after CTCF depletion (**Figure 3C-F**) – spanning a total of around 1.5Mb. For the two loci surveyed, loss of CTCF reduced inter-TAD 3D distances, which became equivalent to intra-TAD distances. This indicates that loss of insulation arises from compacting sequences initially in separate TADs. Intra-TAD FISH distances were unaffected by CTCF depletion, indicating that loss of CTCF does not trigger general chromatin compaction. In the absence of CTCF, linear genomic coordinates becomes a better predictor of 3D distances (Figure S3H) and, consistent with previous boundary-deletion experiments (Ji et al., 2016), TAD boundaries separate further apart in the three-dimensional space of the nucleus (Figure S3I).

Interestingly Hi-C detected fewer intra-TAD contacts (in line with (Zuin et al., 2014)), while FISH did not detect changes in intra-TAD compaction (**Figure 3C-F**). This likely reflects the fact that total Hi-C read number is normalized between samples (so increased inter-TAD signal must be compensated by decreased signal elsewhere) while FISH distances are less resolutive but absolute – a limitation in comparing Hi-C and FISH (Dekker, 2016; Fudenberg and Imakaev, 2016; Giorgetti and Heard, 2016).

### Disruption of Local Insulation Does Not Affect Higher-Order Chromosome Folding

We next sought to investigate to what extent CTCF disruption affects higher-order segregation of active and inactive chromosome domains into A- and B-compartments (Gibcus and Dekker, 2013; Lieberman-Aiden et al., 2009). Inspection of contact maps did not reveal detectable change in the plaid pattern arising from the separate folding of A- and B-types compartments (**Figure 5A**). Close inspection of the compartment signal, computed by ICE (Imakaev et al., 2012), confirms the overall maintenance of compartmentalization and location of transitions between A- and B- compartments (**Figure 5B-C and S5A**). We could only detect a minor but reproducible reduction (˜10%) in the strength of compartmentalization upon CTCF depletion (**Figure S5B**). Scaling of contact frequencies as a function of genomic separation did not change either (**Figure 5D**). This suggests again that long-range (supra-Mb) chromosome architecture is preserved after the loss of CTCF, as well as overall chromatin compaction above the megabase-scale. Factors other than CTCF must therefore control the basal packaging regime of chromatin as well as its segregation in A- and B-compartments.

**Figure 4.**
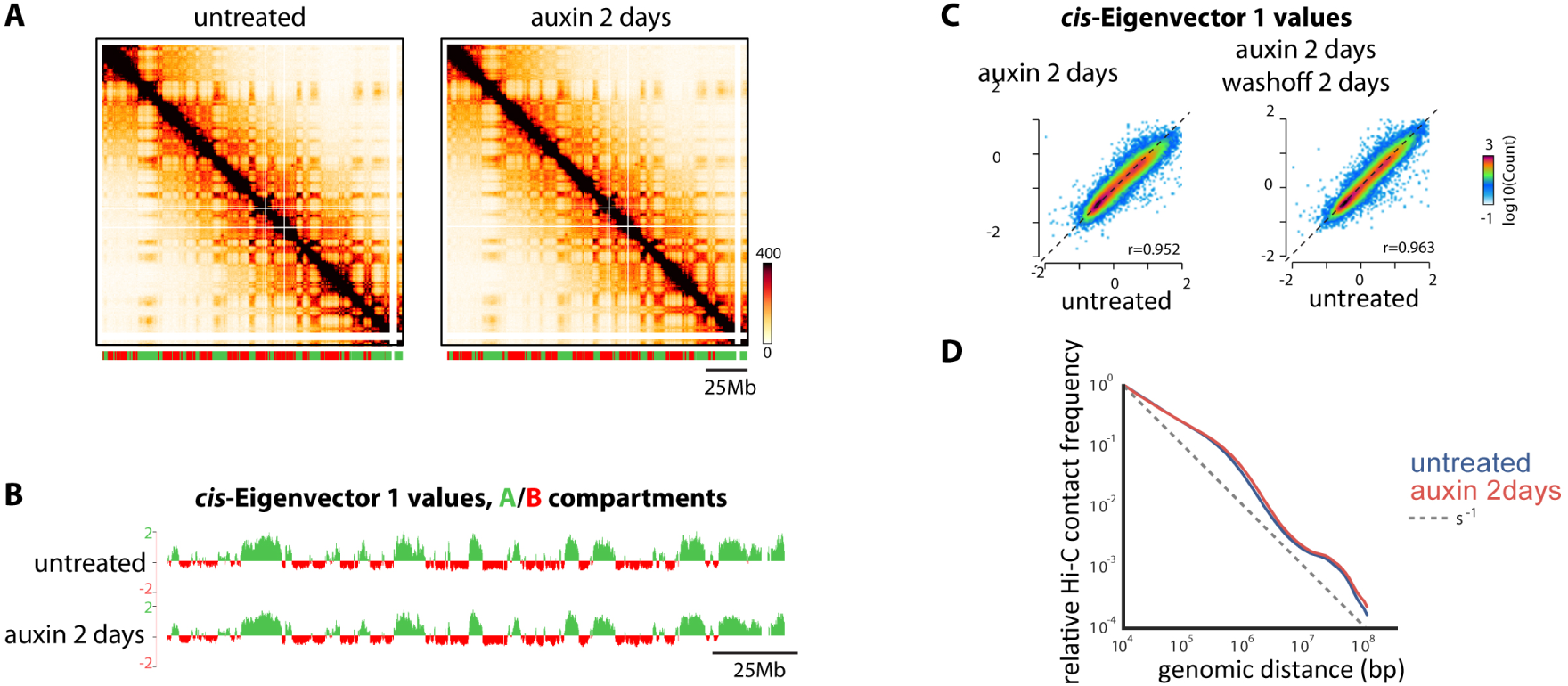
Proper insulation of TADs is not required for higher-order segregation of A and B compartments. (A) Hi-C contact maps at 100kb resolution across entire chromosome 2 highlights maintenance of compartments after CTCF depletion. Bar denotes segments called within A (green) or B (red) compartment using 20kb-*cis* Eigenvector 1. Normalized Hi-C counts are multiplied by 10^5^. (B) Distributions of *cis* Eigenvector 1 values across entire chromosome 2 are remarkably stable to depletion of CTCF. (C) *cis* Eigenvector 1 values are not affected genome-wide by CTCF depletion. **See also Figure S4** (D) Overall scaling of Hi-C contact frequency as a function of genomic distance is not affect by the loss of CTCF, highlighting that CTCF does not affect general chromatin compaction.

**Figure 5.**
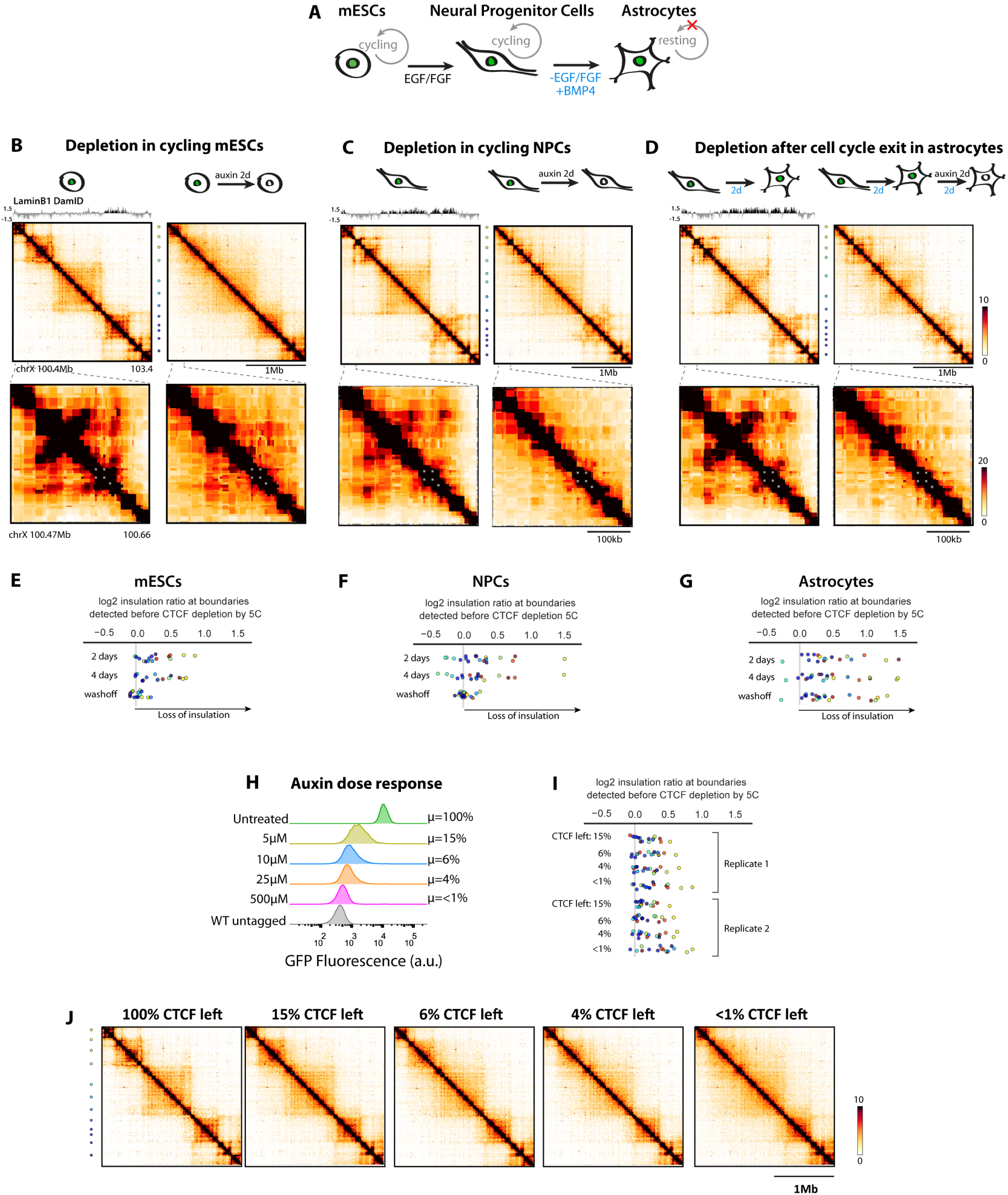
CTCF remains essential for insulation of TADs in resting cells, and acts dose-dependently. (A) Cycling mESCs can be differentiated into cycling NPCs, which can be induced to exit cell cycle by terminal differentiation into astrocytes upon addition of BMP4 and EGF/FGF withdrawal. (B-D) restriction-fragment resolution interpolated 5C heatmaps at the *Xic* reveals that CTCF is required for loops between TAD boundaries and TAD insulation in mESCs, NPCs and resting astrocytes. TAD association to the nuclear periphery identified by LaminB1 DamID from Peric-Hupkes et al. 2010. Color dots denote boundaries identified before CTCF depletion. (E-G) log2 ratio of 100kb insulation scores from depleted versus untreated cells at boundaries identified before depletion, highlighting that a subset of boundaries loose insulation reversibly in mESCs and NPCs but irreversibly in astrocytes. Plots include boundaries probed beyond the region depicted in the heatmaps. (H) Titration of auxin leaves cells with intermediate CTCF levels (I) CTCF-dependent boundaries loose insulation as a function of leftover CTCF levels (J) (J) 5C heatmaps used to calculate insulation scores. **See also Figure S5.**

We next explored if the residual TAD boundaries detected after CTCF depletion (18% of initial boundaries) could be explained by the maintenance of A/B compartmentalization. Out of the 1000 CTCF-less residual boundaries only 103 (10% - 3.1 fold enrichment over chance overlap, **Figure S3E-F**) were associated with a transition between A- and B-compartments. Transcriptional activity (neighboring PolII ChIP-seq peak detected in untreated cells) was detected at 416 of these residual boundaries (41% - 2 fold enrichment over chance overlap). While this is compatible with compartment transition or transcription itself participating in the maintenance of CTCF-independent insulation, either of these features alone is not sufficient to drive CTCF-independent insulation since most boundaries associated with a compartment transitions or PolII are affected by CTCF depletion (Supplementary Table 6). Discrepancies with the reference genome may also account for some of the apparent retention of insulation. This analysis indicates that CTCF is the major driver of insulation of chromatin domains but that other less pervasive processes also participate in insulating mammalian chromosome neighborhoods.

Altogether, local TAD insulation and compartmentalization of mammalian chromosomes therefore appear to be principles that largely rely on different molecular mechanisms.

### Loss of CTCF also Triggers Misfolding in Non-Cycling Cells

To determine if insulation defects triggered by CTCF depletion require passage through DNA replication or mitosis, we differentiated our CTCF-AID mESCs into resting astrocytes (Sofueva et al., 2013). The Tir1 transgene variegated upon differentiation, which we overcame by first converting CTCF-AID mESCs into self-renewing Neural Precursor Cells (NPCs), subcloning NPCs and then selecting clonal lines that retained homogenous CTCF degradation upon auxin treatment. This system provides a unique opportunity to trigger CTCF depletion (by means of auxin addition) before or after exit from the cell cycle (by means of FGF/EGF removal and BMP4 addition, **Figure 5A**) (Peric-Hupkes et al., 2010; Sofueva et al., Peric-Hupkes et al., 2010; Sofueva et al., 2013). 5C at the *Xic* revealed that CTCF depletion triggered loss of contact between CTCF sites and insulation defects at CTCF-dependent boundaries in cycling NPCs as well as resting ACs, whether CTCF was depleted before (**Figure 5B-G**) or after (**Figure S5**) cell-cycle exit. These observations indicate that CTCF controls demarcation into TADs and looping between TAD boundaries through processes that are at play even in non-cycling cells. Folding defects appeared somewhat less pronounced in differentiated cells, and correlate with a large portion of the region surveyed into a lamina associated domain (LAD) – and presumably B compartment. Washing off auxin led to reformation of insulated TADs in mESCs and NPCs but not resting astrocytes, suggesting that passage through the cell cycle might be required for recreating insulation and that factors that cooperate with CTCF *(e.g.* Cohesin metabolism) may behave differently in terminally differentiated cells.

### CTCF depletion Needs to Be Near-Complete to Exhibit the Most Substantial Defects on TAD insulation

Previous studies with RNAi-mediated knock-down of CTCF in human HEK293 cells reported much milder folding defects than those we observed with CTCF-AID mESCs (Zuin et al., 2014). Differences may be attributed to differential sensitivity of mESCs compared to human immortalized cells, or to better depletion efficiency with the degron system than with RNAi, which leaves 10–15% CTCF (Zuin et al.,2014). To formally address this, we treated CTCF-AID mESCs with intermediate doses of auxin and repeated 5C at the *Xic* in the context of various leftover amounts of CTCF, as quantified from fluorescence of the CTCF-AID-eGFP fusion (**Figure 4H**). Insulation defects scaled with the degree of CTCF depletion and samples with around 15% CTCF preserved more insulation than completely depleted cells (with some boundaries more sensitive than others). This highlights that CTCF is very potent at mediating chromatin folding into TADs, acts in a dose-dependent fashion, and must therefore be very efficiently depleted to trigger major defects on chromosome organization.

### CTCF and transcriptional regulation

We then explored how the changes in local genome folding caused by acute CTCF depletion relate to transcriptional misregulation. We performed a time-course RNA-seq experiment in mESCs after 1, 2, or 4 days of auxin treatment (**Figure 6A-B**). The absolute number of differentially expressed genes increased over ten-fold between day 1 (341) and day 4 (4377, over a third of active genes) (**Figure S6A-B**), with around half of the dysregulated being down-regulated and half up-regulated at each timepoint.

**Figure 6.**
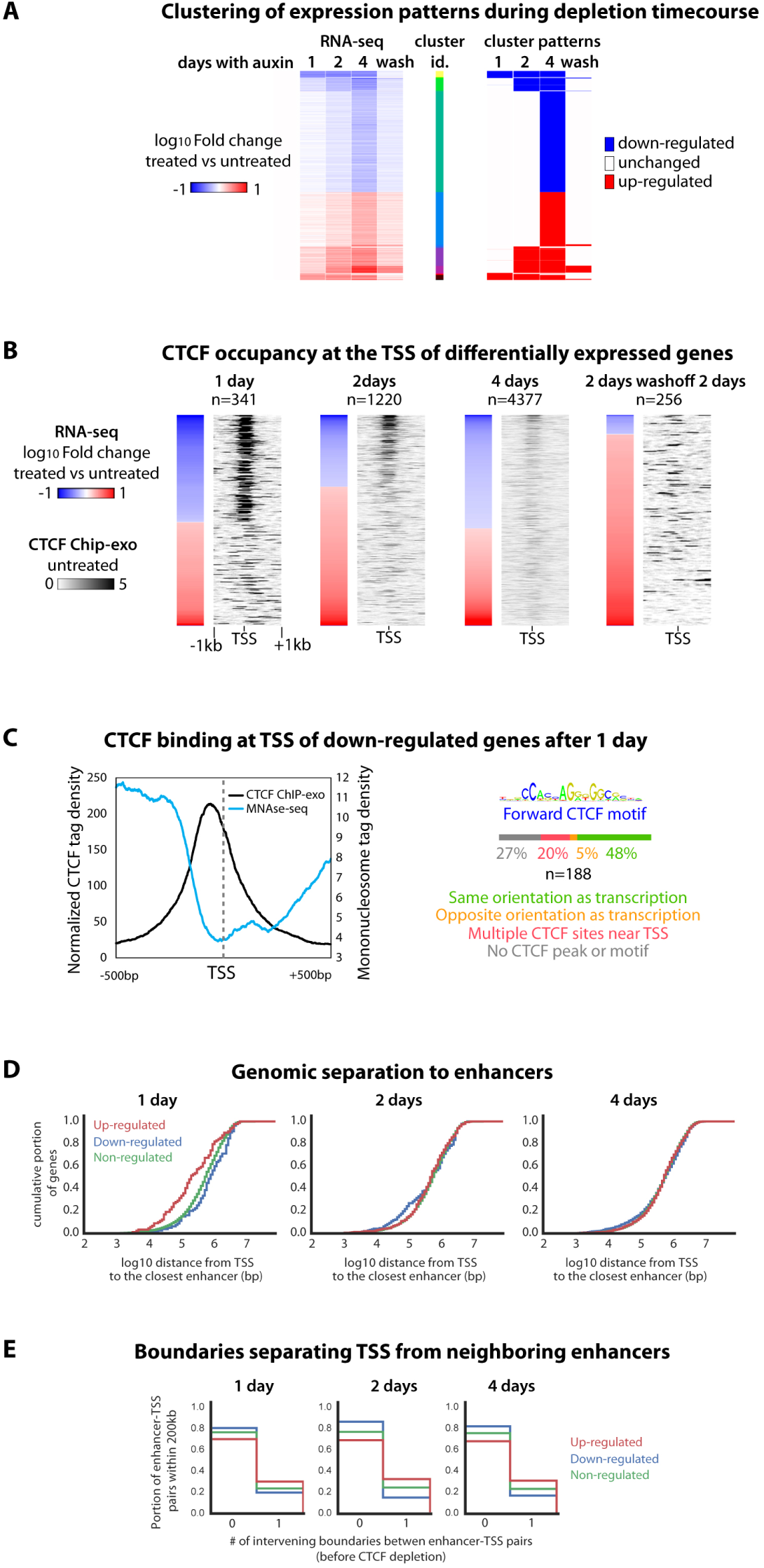
CTCF and transcriptional regulation. (A) RNA-seq timecourse reveals rapid accumulation of transcriptional defects upon CTCF depletion. Fold change compared to untreated cells is shown for genes differentially expressed at one or more time points. Wash denotes 2-day washoff after a 2-day treatment. (B) Alignment with ChIP-exo (from untreated cells) highlights that immediately down-regulated genes had CTCF bound at TSS prior to depletion. Immediately up-regulated genes did not. (C) The CTCF site in the promoters of immediately down‐regulated genes tends to be ^~^60bp upstream of the TSS in direct orientation with transcription, and demarcates the beginning of the nucleosome‐depleted region as previously measured by MNAse‐seq (Teif et al., 2012) (D) Immediately up-regulated genes tend to lie at shorter genomic distance to neighboring enhancers than down-or non-regulated genes. Trend is rapidly lost over time. (E) enhancer-promoter pairs are more likely to be normally interrupted by a TAD boundary for genes that become up-regulated upon CTCF depletion. **See also Figure S6.**

We first focused on down-regulated genes. Integration with CTCF Chip-exo data revealed that over 80% of the early down-regulated genes had CTCF bound within 1kb around the transcription start site (TSS) prior to depletion, as opposed to less than 20% of the up-regulated genes (**Figure 6B**). This trend is diluted with time as the number of differentially expressed genes rises. This highlights that the activity of a subset of CTCF-bound promoters (10% of all CTCF bound TSS) critically relies on CTCF, likely via direct binding. We explored if this activator role may be attributed to CTCF facilitating communication with distal regulatory elements. Out of the 189 genes down-regulated after 1 day of depletion 53 (18%) overlap an anchor for SMC1a HiChIP loops (Mumbach et al., 2016) and 19 (10%) connect to an active regulatory region before treatment, based on H3K27Ac enrichment (Shen et al., 2012). Furthermore, down-regulated genes are not specifically positioned at TAD boundaries. Therefore down-regulation cannot be explained by loss of direct looping between promoters and enhancers. Interestingly, at the promoter of the immediately down-regulated genes CTCF is bound slightly upstream of the TSS (around 60bp) and demarcates the beginning of the nucleosome-depleted region (**Figure 6C**), as visualized by MNAse-seq (Teif et al., 2012). CTCF may therefore promote transcription by preventing promoter occlusion by nucleosomes. Intriguingly, the orientation of the CTCF motif at these TSSs is almost systematically in direct orientation with the direction of transcription (90% of unequivocal sites, **Figure 6C and Supplementary Table 7**). Given the implication of CTCF motif orientation in controlling long-range contacts (Rao et al., 2014; Vietri Rudan et al., 2015) it remains possible that CTCF depletion down-regulates the immediately responsive genes by disrupting enhancer tracking processes that are not associated with accumulation of loops as detected by a peak of Hi-C or HiChIP signal.

We then investigated up-regulated genes, and the possible effect of TAD dissolution on ectopic enhancer targeting. The fact that CTCF does not bind the majority (80%) of TSS of genes up-regulated after 1 day suggests CTCF normally represses them indirectly. Interestingly, immediately up-regulated genes tend to be located genomically closer to active enhancers (**Figure 6D**) or super-enhancers (**Figure S6B**) than down- or non-regulated genes. However, a higher fraction of up-regulated genes normally have a TAD boundary separating them from neighboring enhancers or super-enhancers (<200kb), compared to down-regulated or non-regulated genes (**Figure 6E and S6C**). This suggests that CTCF depletion triggers up-regulation of a subset of genes formerly insulated from neighboring enhancers by a TAD boundary. This observation supports at the genome-wide level the notion that CTCF can mediate enhancer-blocking insulation through the specification of TAD boundaries, in line with previous locus-specific studies (Dowen et al., 2014; Doyle et al., 2014; Lupiáñez et al.; Nora et al., 2012).

Auxin washoff after a 2-day treatment did not completely restore the transcriptome, with most (231 out of 256, 90%) of the differentially expressed genes remaining up-regulated compared to untreated cells (**Figure 6A-B**). Some transcripts showed a trend downward their initial values while others kept rising (**Supplementary Table 3**), suggesting that for a small subset of genes transient loss of CTCF depletion can trigger transcriptional changes that rapidly become irreversible, suggesting they are involved in a positive feedback mechanism.

### CTCF Binding Is Not a Direct Impediment to H3K27me3 Spreading in mESCs

It has been suggested that CTCF may confer chromatin barrier activity by opposing the spreading of facultative heterochromatin, thereby demarcating active and inactive chromatin domains (Cuddapah et al., 2009; Dowen et al., 2014) and insulating against position effects (Essafi et al., 2011; Witcher and Emerson, 2009). This role has been debated (Bender et al., 2006; Huang et al., 2007; Ong and Corces, 2014; Recillas-Targa et al., 2002; Splinter, 2006)‥

As reported in human cells (Cuddapah et al., 2009), we found that a subset of CTCF sites mark the boundaries of H3K27me3-rich regions in mESCs (^~^7% of CTCF sites, **Figure 7A**). However CTCF depletion did not trigger spreading of H3K27me3 as measured by ChIP-seq, even after 4 days (3-4 cell divisions **Figure S1B**). This argues against a direct chromatin barrier function for CTCF (**Figure 7B-C**). Changes were restricted to a very local gain of H3K27me3 signal at the formerly occupied CTCF site (**Figure 7B and S7A**), possibly due to nucleosomes becoming able to occupy the formerly bound CTCF site (Fu et al., 2008; Wiechens et al., 2016). On a more global scale, we observed a slight but significant decrease in overall H3K27me3 levels (**Figure S7B**). These changes are likely indirect effects as they are not restricted to the vicinity of CTCF sites; they may be accounted for by the 2-fold transcriptional down-regulation of the essential PRC2 component EED, especially after 4 days of depletion (Supplementary Table 3).

**Figure 7.**
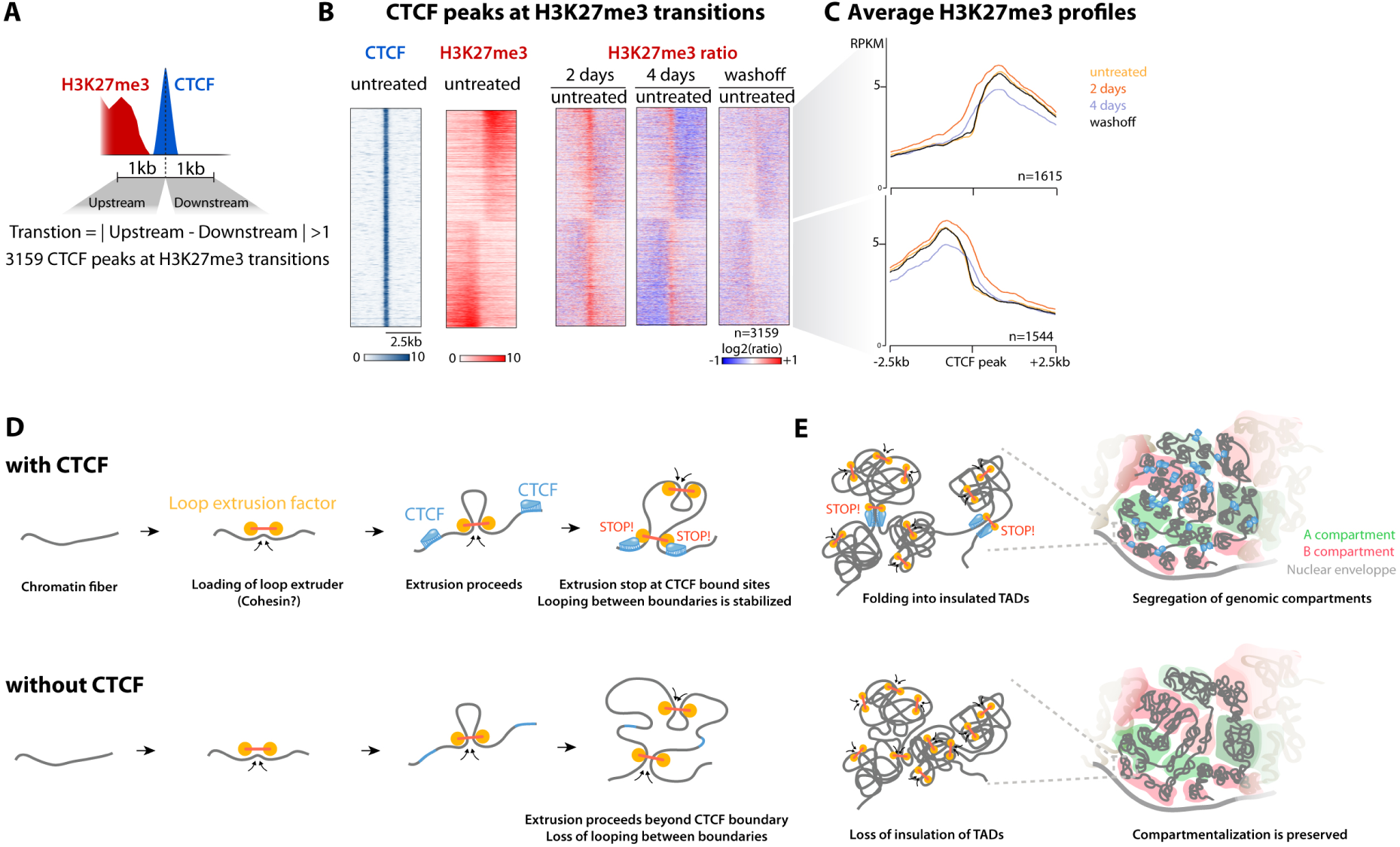
CTCF does not constrain H3K27me3 spreading in mESCs and summary model. (A) A subset of CTCF binding sites mark transitions in H3K27me3 patterns (B-C) CTCF depletion does H3K27me3 spreading beyond the formerly bound CTCF site itself (center). **See also Figure S7.** (D) Our observations are consistent with TAD formation by loop extrusion, and establish CTCF as the major factor defining domain boundaries genome-wide. (E) Statistical representation of TAD disruption and compartment preservation upon loss of CTCF.

Altogether our results demonstrate that the role of CTCF in genome organization is local, in controlling the accumulation of chromatin loops between TAD boundaries and physically insulating these domains from each other. In the absence of CTCF neighboring TADs merge, with consequences on transcriptional regulation. Overall chromosome compaction and organization are however not affected. Other factors than CTCF must therefore be responsible for general chromatin packaging and compartmentalization.

## DISCUSSION

Using a cellular system enabling acute, reversible and near-complete loss of CTCF, we have elucidated the critical and dose-dependent roles of this enigmatic transcription factor in regulating 3D chromatin organization. Beyond establishing the central importance of CTCF in insulation of TADs, this system has enabled to address fundamental questions about the causal relationships between the different levels of genome organization, transcription, and large-scale chromatin states. Our findings are consistent with spatial compartmentalization of mammalian genomes being a high-order emerging property that relies on molecular mechanisms that are distinct from those controlling the local insulation of chromosome neighborhoods.

### CTCF Is Necessary for TAD Insulation and Loops between Boundaries

CTCF depletion concomitantly disrupts loops between TAD boundaries and insulation of neighboring TADs. This substantiates the notion that these two aspects are molecularly coupled (Giorgetti et al., 2014). Our observations are compatible with mechanistic models where domain-wide enrichment of chromosomal contact is the result of a process that accumulates chromatin loops between CTCF-bound boundary elements (Fudenberg et al., 2016; Sanborn et al., 2015) (**Figure 7D**).

### Pervasive Loss of Insulation upon CTCF Depletion Ascertains the Central Importance of Boundary elements

Two classes of models have been proposed to account for the physical insulation of chromosome domains: boundary effects and block copolymer incompatibility (Imakaev et al., 2015). Our data support the first model by demonstrating that CTCF confers the physical insulating function of TAD boundaries and corroborating, at the genome-wide level, observations from previous locus-specific boundary deletion experiments (Darrow et al., 2016; Giorgetti et al., 2016; Hnisz et al., 2016; Lupiáñez et al.; Nora et al., 2012; Sanborn et al., 2015). The restoration of boundary activity and chromatin folding into TADs upon reinstatement of CTCF demonstrates that CTCF carries sufficient instructive power to enable accurate TAD folding. How this is achieved mechanistically will be the focus of future research.

Block copolymer models posit that intrinsic interaction incompatibility between neighboring TADs is the determinant for their insulation (Chiariello et al., 2016). This may explain the existence of CTCF-independent boundaries at some transitions between A and B compartments. However these CTCF-independent boundaries only represent a small fraction of all boundaries (less than 18%). Furthermore, most of boundaries at compartment transitions lose insulation upon CTCF depletion (**Supplementary figure S4C**), arguing that compartmentalization in itself does not strongly contribute to local insulation of TADs. Insulation through block co-polymer incompatibility may be more relevant in other biological contexts where chromatin states are a better predictor of segmental packaging into TADs, such as in *Drosophila melanogaster* (El-Sharnouby et al., 2016; Jost et al., 2014; Ulianov et al., 2015).

### CTCF-Depletion Reveals Local Insulation and A/B Compartmentalization Are Molecularly Independent Principles of Mammalian Genome Folding

Long-range chromosome folding (above the megabase scale) appears remarkably resistant to CTCF depletion, in the context of dramatic changes at the sub-megabase scale. We conclude that proper packaging of chromatin into TADs is not a prerequisite for the segregation of A and B compartments. It is possible that the precise boundaries of the chromosomal segments belonging to the same type of compartment are slightly altered at scales below what can be detected with our current 20-kb resolution.

This finding corroborates recently documented cases in which TAD folding and compartmentalization are uncoupled, such as the *Drosophila* polytene chromosomes that insulate TADs without compartmentalizing them (Eagen et al., 2015).

Our observations are consistent with the proposed mechanisms of TAD formation by intra-TAD loop extrusion and support the idea that CTCF is a major blocking factor to the processivity of extrusion (Fudenberg et al., 2016; Sanborn et al., 2015). Notably, the extrusion model accurately describes mammalian chromosome folding at the submegabase scale but does not account for the segregation of genomic compartments. The molecular drivers of CTCF-independent higher-order compartmentalization remain to be defined. The characteristic epigenomic signature of each (sub)compartment suggests that chromatin states or chromatin-binding proteins may have a role (Kind et al., 2015; Lieberman-Aiden et al., 2009; Rao et al., 2014). Indeed, a fraction of H3K27me3 marked loci connect preferentially with each other (Joshi et al., 2015; Vieux-Rochas et al., 2015) in a fashion that depends on the Polycomb machinery (Schoenfelder et al., 2015). Furthermore, artificially targeting heterochromatin proteins, such as Ezh2, can induce domain-wide compartment switching (Wijchers et al., 2016). Such chromatin-based mechanisms of compartmentalization do not appear to affect local insulation as proper folding into TAD occurs in the absence of H3K27me3 or H3K9me2 (Nora et al., 2012), again pointing to mechanistic differences between formation of insulated TADs and higher order A- and B- compartments.

The stability of chromatin states upon CTCF depletion (at least for H3K27me3) may explain the maintenance of long-range (supra-megabase) chromosome folding. This suggests that organisms or cell types that lack CTCF-mediated insulation mechanisms can instead rely on compartmentalization to spatially organize their genome.

### CTCF Does Not Directly Constrain the Spread of H3K27me3 but May Still Define Chromatin Domains

Our observation that H3K27me3 patterns remain largely unaltered challenges the notion that CTCF binding acts as a direct impediment to heterochromatin spreading. This is consistent with the absence of H3K27me3 spreading after serial genetic deletions of the *HoxD* locus, removing large segments, including CTCF sites (Schorderet et al., 2013). Our observations in undifferentiated ES cells do not, however, address the role of CTCF binding in defining the genomic segments that can undergo domain-wide chromatin state transitions during cell differentiation, which were initially found to align with TAD boundaries (Nora et al., 2012). Interestingly, deleting single CTCF sites within the *HoxA* cluster enables ectopic developmental activation of genes across the former boundary, consistent with ectopic enhancer targeting, but again does not lead to H3K27me3 spreading (Narendra et al., 2015). Altogether current data support that CTCF mediates enhancer-blocker activity, through its ability to mediate insulation and segmental folding into TADs, and is not a direct impediment to heterochromatin spreading.

### TAD Insulation and Gene Regulation

The pervasiveness of the chromosome folding defects we observed upon CTCF depletion contrasted with the rather limited immediate transcriptional defects measured by RNA-seq. It is difficult to interpret prolonged depletion as secondary effects can rapidly become confounding. Our data highlight that exposure of a promoter to new enhancers has an initially mild and context-specific impact on transcriptional activity. This suggests that hijacking of *cis*-regulatory elements caused by altered insulation may require time to manifest pervasively, and that ectopic contact between enhancers and promoters is not in itself sufficient to predict the initial extent of transcriptional defects. Additional specificity or compatibility factors must contribute to how promoters respond after ectopic exposure to enhancers (van Arensbergen et al., 2014; Zabidi et al., 2015).

Interestingly, we did not observe immediate coordinated TAD-wide transcriptional changes. This may appear at odds with previous reports of TAD-wide coordination of transcription dynamics upon deletion of a TAD boundary or during early response to signaling (Le Dily et al., 2014; Narendra et al., 2015; Nora et al., 2012). The timing needed for transcriptional defects to manifest themselves may explain this apparent discrepancy. Boundary disruption experiments are typically analyzed at the steady-state in cells or organisms long after the rearrangement has been induced, after cells have adapted (Dowen et al., 2014; Flavahan et al., 2016; Franke et al., 2016; Hnisz et al., 2016; Taberlay et al., 2016; Tsujimura et al., 2014). On the other hand acute degradation of CTCF provides the opportunity to monitor immediate effects, but is also expected to trigger a wide range of effects, where direct but slowly manifesting effect will be obscured by indirect but rapid secondary effects.

In summary our findings reveal that distinct molecular mechanisms orchestrate mammalian chromosome packaging, segmentation into TADs and higher-order compartmentalization. Our study also demonstrates the feasibility of coupling the AID system with genome editing to study endogenous mammalian transcription factors in stem cells. We anticipate its further use will be instrumental in interrogating causal relationships in epigenomics, especially when studying essential factors.

## AUTHOR CONTRIBUTIONS

E.P.N. conceived and designed the study with input from B.G.B; E.P.N. engineered and cultured cell lines, performed 5C, ChIP-exo, ChIP-seq, RNA-seq and FISH with help from A.U., and analyzed data. A.-L.V. and J.B., performed Hi-C in the lab of J.D and pre-processed Hi-C and 5C data. A.G. and N.A. performed computational analyses of Hi-C in the lab of L.A.M. E.P.N. wrote the manuscript with B.G.B and with input from all authors.

## ACKNOWLEDGEMENTS

We apologize for not citing numerous relevant studies due to space constraints. We thank Daphné Dambournet, Baptiste Roelens and Matthias Merkenschlager for discussions; Casey Gifford for help with sequencing; Elizabeth Blackburn for access to microscopy resources; Sean Thomas, Alex Williams and the Gladstone Bioinformatics core; Gary Howard for editorial assistance; Edith Heard and Geoffrey Fudenberg for critical comments on the manuscript. This work was supported by the EMBO (ALTF523-2013) and HSFP (E.P.N.); the National Institutes of Health/National Heart, Lung, and Blood Institute (Bench to Bassinet Program U01HL098179), the Gladstone Institutes, and William H Younger, Jr. (B.G.B); the UCSF-Gladstone Institute of Virology & Immunology Center for AIDS Research (CFAR) NIH P30 AI027763 (Gladstone flow cytometry core); the National Human Genome Research Institute (R01 HG003143, U54 HG007010, U01 HG007910), the National Cancer Institute (U54 CA193419), the NIH Common Fund (U54 DK107980, U01 DA 040588), the National Institute of General Medical Sciences (R01 GM 112720), and the National Institute of Allergy and Infectious Diseases (U01 R01 AI 117839). J.D. is an investigator of the Howard Hughes Medical Institute. B.G.B. is a co-founder of Tenaya Therapeutics.

## METHODS

### Key resources Table

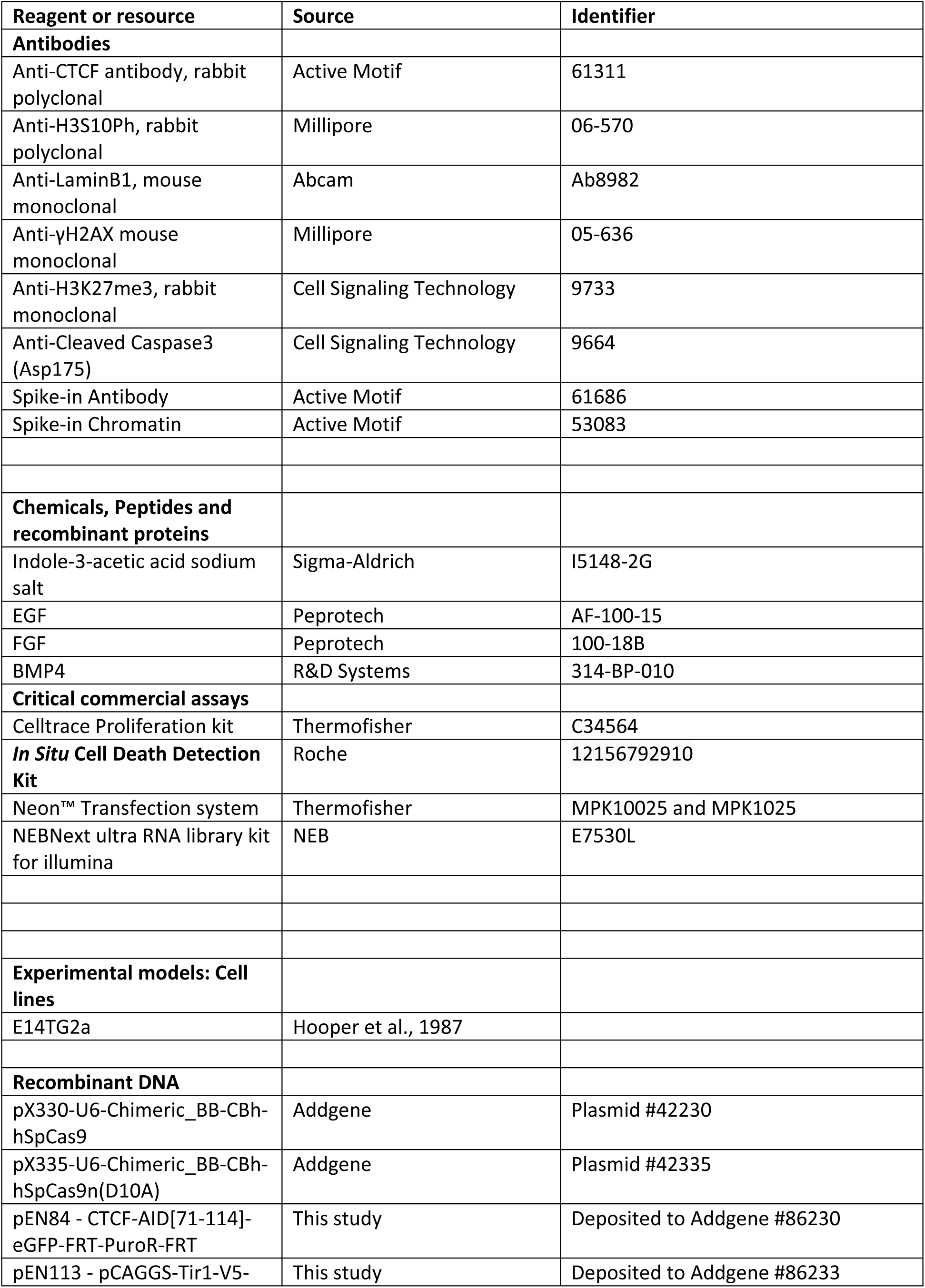

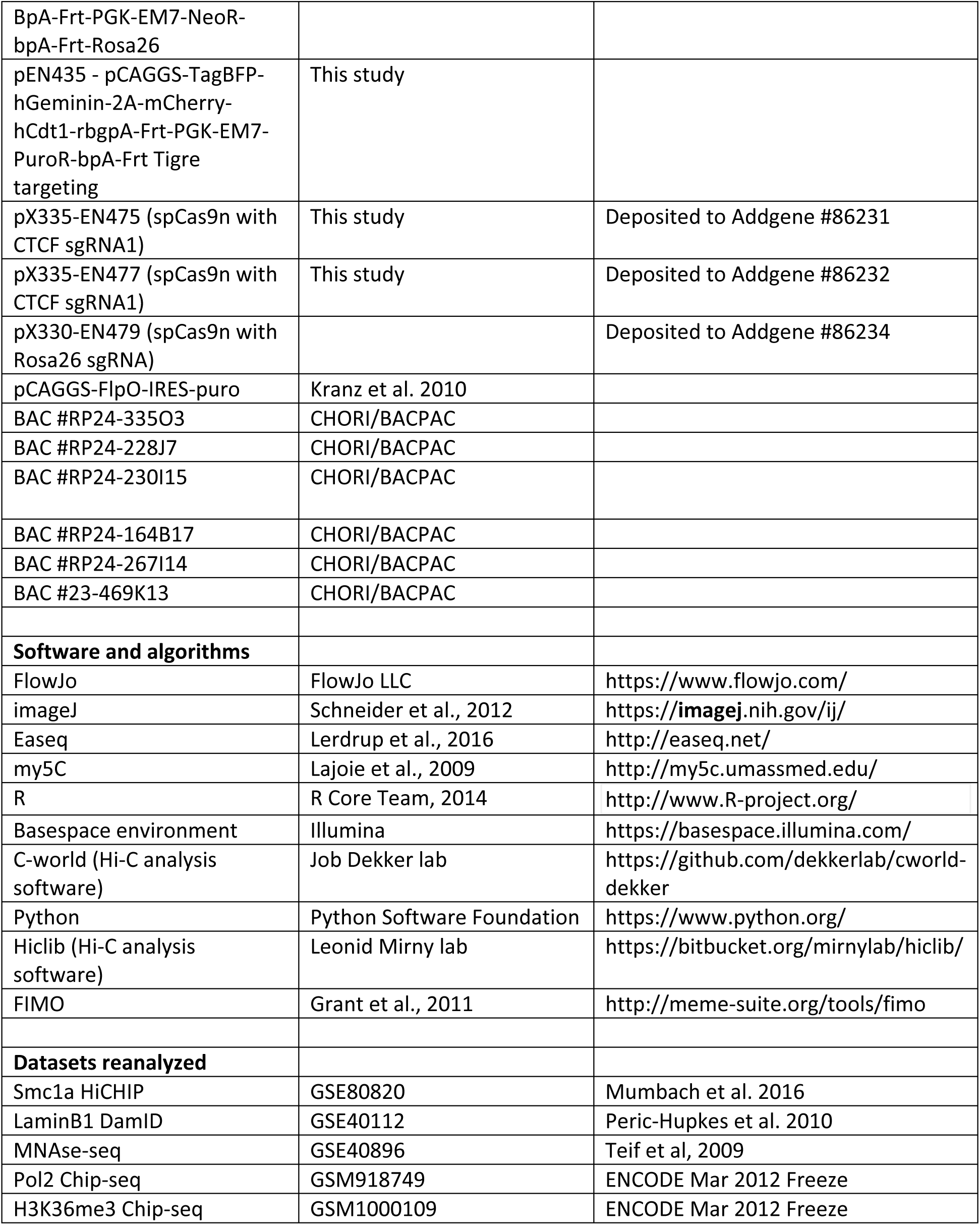

### Contact for Reagent and Resource Sharing

Further information and requests for reagents may be directed to and will be fulfilled by the corresponding authors Benoit Bruneau and Elphège Nora.

## Experimental Model and Subject Details

### Mouse Embryonic Stem cells

E14Tg2a (karyotype 19, XY; 129/Ola isogenic background) and subclones were cultured in DMEM+Glutamax (ThermoFisher cat 10566-016) supplemented with 15% Fetal Bovine Serum (ThermoFisher SH30071.03), 550μM b-mercaptoethanol (ThermoFisher 21985-023), 1mM Sodium Pyruvate (ThermoFisher 11360-070), 1X non-essential amino-acids (ThermoFisher 11140-50) and 10^4^U of Leukemia inhibitory factor (Millipore ESG1107). Cells were maintained at a density of 0.2-1.5×10^5^ cells / cm^2^ by passaging using TrypLE (12563011) every 24-48h on 0.1% gelatin-coated dishes (Millipore cat ES-006-B) at 37°C and 7% CO2. Medium was changed daily when cells were not passaged. Cells were checked for mycoplasma infection every 3-4 months and tested negative.

### Neural progenitor cells and astrocytes

CTCF-AID mESCs were seeded at around 0.1 million cells in a 75cm^2^ gelatinized dish in mESC medium. The following day cells were rinsed twice in 1X PBS and switched to NDiff227 differentiation medium (Stem Cells Inc.) and changed daily. After 7 days cells were detached using TryplE and seeded on nongelatinized bacterial dishes for suspension culture at 3 million cells per 75cm^2^ and cultured in NDiff227 containing 10ng/mL EGF and FGF (Peprotech). After 3 days floating aggregates were seeded on gelatinized dishes. After 2-4 days cells were dissociated using Accutase and passaged twice on gelatinized dishes in NDiff227+EGF+FGF. In order to overcome variable silencing of the Tir1 transgene the CTCF-AID NPCs were subcloned by limiting dilution and NPC colonies were manually picked after 10-15 days and expanded in NDiff227+EGF+FGF. For differentiation into quiescent astrocytes adherent NPC cultures were washed twice with NDiff227 and cultured for at least 48h with NDiff227+ 10ng/mL BMP4 (R&D Systems). The CTCF-AID NPC subclones did not survive freeze and thawing.

For induction of the auxin-inducible degron indole-3-acetic acid (IAA, chemical analog of auxin) was added in the medium at 500μM from a 1000X stock diluted in sterile water. Stocks were kept at 4°C up to 4 weeks or −20°C for long term storage.

## Method Details

### Plasmid Construction

We used the smallest functional truncation of the AID tag (AID*, 44 aino-acids), initially developed in yeast (Morawska and Ulrich, 2013), and observed equivalent CTCF depletion efficiency, compared to the original full-length 231 amino-acid tag (Nishimura et al., 2009) (data not shown).

The CTCF-AID-EGFP targeting vector (pEN84) was assembled by serial modification of the base vector pFNF (Addgene #22687) using Gibson assembly with the following templates: the minimal functional AID tag (aa 71-114) described by (Morawska and Ulrich, 2013) was PCR amplified from pAID (Nishimura et al., 2009); homology arms to the last exon of *Ctcf* were PCR amplified from E14Tg2A genomic DNA (1kb each); the N-acteyl-transferase (PAC/PuroR) was PCR amplified from pLox-STOP-Lox TOPO (Addgene # 11854), the eGFP cDNA was PCR amplified from pTRE2-2A-eGFP (Kind gift from Kevin Monahan and Stavros Lomvardas).

The Tir1 expression vector (pEN113) was assembled by serial modification of the base vector pFNF (Addgene #22687) using Gibson assembly with the following templates: CAGGS promoter was subcloned from pCAGEN (Addgene #11160), the Oryza Sativa Tir1 cDNA was PCR amplified from a synthetic mammalian codon-optimized vector (kind gift from Daphné Dambournet and David Drubin); homology arms to the *Rosa26* locus were PCR amplified from E14Tg2A genomic DNA (1kb each).

The BFP/mCherry FUCCI reporter (pEN435) was assembled by serial modification of the base vector vector pFNF (Addgene #22687) using Gibson assembly with the following templates: hGeminin and mCherry-Cdt1 were PCR amplified from pRetroX-S2G2M and pRetroX-G1-Red (Clonetech); tagBFP cDNA from pHR-Tet3G-2A-BFP (Kind gift from Stanley Qi); CAGGS promoter and puroR are of the same source as pEN113; homology arms to the *Tigre* locus (Zeng et al., 2008) were PCR amplified from E14Tg2A genomic DNA (1kb each).

Maps of the targeting constructs in the Genbank format are available on Addgene.

sgRNAs were cloned by annealing pairs of oligos either in pX330 (Addgene #42230) for single Cas9 nuclease or pX335 (Addgene #42335) for dual Cas9 nickase strategies, following the protocol described in (Cong et al., 2013). *Ctcf*-targeting sgRNAs were cloned in pX335 (dual nickase) by annealing oligos caccgATCACCGGTCCATCATGCTG and aaacCAGCATGATGGACCGGTGATc for the first sgRNA and caccgCTGGGGCCTTGCTCGGCACC and aaacGGTGCCGAGCAAGGCCCCAGc for the second sgRNA. *Rosa26* sgRNAs were cloned in pX335 (dual nickase) by annealing oligos caccgTGGGCGGGAGTCTTCTGGGC and aaacGCCCAGAAGACTCCCGCCCAc for the first sgRNA and caccgACTGGAGTTGCAGATCACGA with aaacTCGTGATCTGCAACTCCAGTc for the second sgRNA (note these sgRNAs underperformed, see below). The *Tigre*-targeting sgRNA was cloned into pX330 (single nuclease) by annealing caccgACTGCCATAACACCTAACTT and aaacAAGTTAGGTGTTATGGCAGTc.

### Gene targeting

For transfection plasmids were prepared using the Nucleobond Maxi kit (Macherey Nagel) followed by ethanol precipitation. Constructs were not linearized.

To knock in the AID-eGFP cassette at the N-terinus of CTCF E14Tg2a passage 19 were transfected by microporation using the Neon system (Thermofisher) using a 100μL tip with 1 million cells at 1400V, 10ms and 3 pulses. 2μg of each Ctcf-targeting sgRNA and 20μg of targeting construct (pEN84) was used. After electroporation cells were seeded in a 9cm^2^ well and left to recover for 48h, at which stage around 10% of the cells show nuclear GFP fluorescence. Puromycine was then added to the media at 1μg/mL and cells were selected as a heterogenous pool of homozygous and heterozygous cells for around 10 days, at which stage over 95% of the cells showed nuclear GFP fluorescence. Cells were then transfected with the Neon system using a 10μL tip and 0.1 million cells with 250ng of a flippase-expressing plasmid (pCAGGS-FlpO-IRES-puro) in order to trigger FRT recombination and excision of the puromycine selection cassette. After electroporation cells were seeded in a 9cm^2^ well and left to recover for 48h and transferred into a 78cm^2^ petri dish from whish two serial 1:10 dilution were seeded in an additional two dishes. After 7-8 days of culture without antibiotic selection single colonies were manually picked, transferred into a 96-well plate, dissociated and re-plated. Clones were then genotyped by PCR for homozygous insertion of AID-eGFP and excision of the puro cassette. Over 95% of cells had one knock-in allele, of which 20% were homozygous. Half of the clones were found to have undergone FlpO-mediated recombination. When homozygous both alleles always underwent recombination.

To knock in the Tir1-expressing cassette one homozygous CTCF-AID-eGFP clone was transfected as described above using a 100μL tip format and pEN114 as the targeting construct. After a 48h recovery cells were subcloned and grown for 7 days in the presence of 200μg/μL Geneticin until single colonies could be picked. We noticed that only a handful of resistant clones were recovered, suggesting sub-optimal targeting – either because of the sgRNA or the targeting construct. Clonal lines were assessed for their ability to undergo auxin-mediated degradation of CTCF-AID-eGFP. We selected the clone with the fewest GFP-positive cells (<1%) after 24h of auxin treatment. This clone was then used for transient transfection of pCAGGS-FlpO-IRES-puro as described above to yield the CTCF-AID-eGFP, Tir1 line with which we conducted experiments presented in this manuscript (puromycine and neomycine sensitive). Rosa26 PCR genotyping revealed this clone had undergone random insertion of the Tir1 cassette.

Robust expression of the Tir1 transgene was absolutely critical to mediate auxin responsiveness. Indeed our CTCF-AID lines down-regulated Tir1 during differentiation, even when targeted at *Rosa26,* leading to variegation of auxin response and limiting our analyses in committed cells that can be subcloned, such as neural progenitors. Further improvements in transgenesis will be necessary to enable reliable use of the AID system in both stem cells and their differentiated derivatives.

### Crystal Violet staining

Limiting dilutions of mESCs were plated and grown for 14 days, after which they were rinsed with PBS and fixed/stained with 1% Formaldehyde 1% Methanol in PBS 0.05%w/v Crystal violet for 20 min. Plates were thoroughly rinsed with tap water and air dried.

### Flow Cytometry

mESCs were dissociated with TryplE, resuspended in culture medium, spun and resuspended in 4% FBS-PBS before live flow cytometry on a MACSQuant instrument (Miltenyibiotec). Dissociation, wahs and Flow buffers were supplemented with auxin, when appropriate, to avoid re-expression of the CTCF-AID-eGFP fusion. Analysis was performed using the Flowjo sowftware.

### CellTrace (CFSE) proliferation assay

Dissociated mESCs were labeled with CellTrace Violet dye (ThermoFisher) for 30min in PBS and washed following manufacturer’s recommendations. Initial staining was measured by flow cytometry after 30 min, cells were plated and eventually treated with auxin. Remaining fluorescence was then measured daily for up to 4 days after cell dissociation by flow cytometry.

### Western blots

mESCs were dissociated, resuspended in culture medium, pelleted, washed in PBS, pelleted again and kept at −80°C. 15-20 million cells were used to prepare nuclear extracts. Cell pellets were resuspended in 10mM Hepes pH 7.9, 2.5mM MgCl2, 0.25M sucrose, 0.1% NP40, 1mM DTT, 1X HALT protease inhibitors (ThermoFisher) and swell for 10 min on ice. After centrifugation at 500g nuclei were resuspended in on ice in (25mM Hepes pH 7.9, 1.5mM MgCl2, 700 mM NaCl, 0.5mM DTT, 0.1 mM EDTA, 20% glycerol, 1mM DTT, sonicated and centrifuged at 18,000g at 4°C for 10 minutes. Protein concentration from supernatants were measured using the Pierce Coomassie Plus assay kit (Thermofisher). For CTCF 10μg of nuclear extracts were loaded per lane while for histones 3μg were used. Samples where mixed with Laemmli buffer and 0.025% b-mercaptoethanol final, run on a 4-12% polyacrylamide TGX gel (Biorad). Transfer onto PVDF membranes was performed using the iBlot system (Thermofisher) Program 0 for 8 minutes. Membranes were incubated at least 30 minutes with Odyssey blocking buffer (Li-cor) prior to antibody incubation overnight at 4°C, following manufacturer's recommended dilutions and supplementing with 0.1% Tween-20 and 0.01%SDS. Membranes were washed five times 5minutes in PBS-0.1% Tween-20 at room temperature, incubated with secondaries antibodies (Goat Anti-Rabbit 680RD and Donkey Anti-Mouse 800CW (Licor), 1:10,000) in Odyssey blocking buffer with 0.1% Tween-20 and 0.01% SDS 1h at room temperature, washed 5 times and analyzed on a Li-cor imaging system. Pannels were mounted using imageJ preserving linearity.

### Cell-cycle Analysis by propidium iodide staining

mESCs were dissociated, resuspended in culture medium, pelleted, washed in PBS, resuspended in ice-cold PBS at 2 million cells / mL. 9mL of 70% ethanol was then added drop-wise while mixing and cells were stored overnight at −20°C. Cells were pelleted at 200g 10 min at 4°C, washed with PBS, pelleted again and resuspended in 300μL of 0.1% Triton X-100 in PBS supplemented with 20μg/mL Propidium iodide and 0.2mg/mL RNAse A. After 30min incubation at 37°C cells were transferred on ice and used directly for flow cytometry.

### Immunofluorescence

mESCs were grown on glass-coverslips, fixed with 3% formaldehyde in 1XPBS for 10’ at room temperature. Permeabilization was carried out in 0.5% Triton followed by blocking with 1% BSA diluted in 1X PBS (Gemini cat 700-110) for 15min at room temperature. Primary antibody (1/250) incubation was performed at room temperature for 45min, followed by three 5-minute washes in 1X PBS, secondary antibody (1/10.000) incubation, three 5-minute washes in 1X PBS, counter-staining with DAPI and mounting in 90% glycerol – 0.1X PB – 0.1% p-phenylenediamine pH9.

### 3D-DNA FISH

Procedure was carried out exactly as described in Nora et al. 2012. Probes were prepared by nick translation from following Bacterial Artificial Chromosomes obtained from CHORI/BACPAC:

Tbx5 locus: RP24-164B17, RP24-267I14, RP23-469K13.

Prdm14 locus: RP24-335O3, RP24-228J7, RP24-230I15.

### Microscopy

Images were acquired on a DeltaVision widefield system (GE Healthcare) using a 100X objective and no binning. Images were deconvolved directly with the Softworks software.

### ChIP-seq

For fixation mESCs were dissociated using TrypLE and resuspended in 10% FBS in PBS, counted and adjusted to 1 million cells per mL. Formaldehyde was then added to 1% final followed by 10 minute incubation at room temperature. Quenching was performed by adding 2.5M Glycine-PBS to 0.125M final followed by 5 min incubation at room temperature, 15 minute incubation at 4°C, centrifugation at 200g 5 minutes at 4°C, resuspended with 0.125M Glycine in PBS at 10 million cells per mL, aliquoted, spun at at 200g 5 minutes at 4°C and snap frozen on dry ice.

Fixed cells were thawed on ice, resuspended in ice cold 5mM PIPES pH 7.5, 85mM Kcl, 1% NP-40 and 1X HALT protease inhibitor, counted and readjusted to obtain 10 million cells total exactly, incubated on ice 15 min, centrifuged at 500g 5 min at 4°c, resuspended in 1mL 50mM Tris-HCl pH8, 10mM EDTA pH8, 1% SDS and 1X HALT protease inhibitor, transferred to a MilliTube (Covaris). Chromatin was sheared on a Covaris S2 sonicator for 7 minutes at 5% duty cycle, intensity 8, 200 cycles per burst in a waterbath maintained at 4°C, using 1 min sonication – 30 sec rest, resulting in 200-800bp fragments. Samples were clarified by centrifugation at 18,000g at 4°C for 10 min. Supernatents were transferred to 15mL conicals and 40ng of spike-in *Drosophila* chromatin (Active Motif) was added. 10% of the mixture was saved as input and the rest was diluted to 5mL with ice-cold 50mM Tris-Hcl pH 7.4, 150mM NacCl, 1% NP-40, 0.25% Sodium Deoxycholate, 1mM EDTA, 1X protease inhibitor. 10μg of anti-CTCF together with 4μg spike-in antibody (anti-H2Av, Active motif) or anti-H3K27me3 antibody together with 4μg spike-in antibody (Active motif) was added alongside with 40μL prewashed protein G Dynabeads (ThermoFisher) followed by overnight incubation at 4°C on a rotator. Beads were then collected on a magnetic rack and washed twice with 1mL cold 50mM Tris-Hcl pH 7.4, 150mM NacCl, 1% NP-40, 0.25% Sodium Deoxycholate, 1mM EDTA, twice with 1mL cold 100mM Tris-HCl pH9, 500mM LiCl, 1% NP-40, 1% Sodium deoxycholate and once with 1mL cold 100mM Tris-HCl pH9, 150mM Nacl, 500mM LiCl, 1% NP-40, 1% Sodium deoxycholate. Beads were then eluted with 100μL 50mM NaHCO3 1% SDS and heated at 65°C 30min with shaking. Input sample volumes were adjusted to 100μL with the same buffer. Eluates and inputs were supplemented with 10μg RNAse A and incubated 30 min at 30°C, then 20μg Proteinase K and 12μL of 5M NaCl were added followed by overnight incubation at 65°C. Samples were then purified using 1.8X Agencourt AMPure XP beads (Beckman-coulter) and eluted in 30μL Tris-HCl.

The entire Chip material or 50ng of the input DNA were used to construct Illumina sequencing libraries. End repair was performed in 100μL with 400μM dNTP, 15U T4 DNA polymerase (NEB), 5U Klenow large fragment DNA polymerase (NEB) and 50U T4 PNK (NEB) in 1X T4 ligase buffer (NEB), at room temperature 30min, followed by 1X AMPure purification. Entire eluate was used for A-tailing in a 50μL reaction with 1mM dATP and 15U Klenow 3’->5’ exo minus in 1X NEB buffer 2 followed by 1X AMPure purification. Entire eluate was used for adapter ligation in 50μL with 6,000U T4 ligase (NEB) and 20nM annealed and indexed adapters in 1X T4 ligase buffer (NEB) at room temperature for 2 hours, followed by 0.8X AMPure purification. Adapters were prepared by annealing following HPLC purified oligos: 5’-AATGATACGGCGACCACCGAGATCTACACTCTTTCCCTACACGACGCTCTTCCGATC*T and 5’Phos-GATCGGAAGAGCACACGTCTGAACTCCAGTCACNNNNNNATCTCGTATGCCGTCTTCTGCTTG T where * represents a phosphothiorate bond and NNNNNN is a Truseq index sequence.. The entire eluate was then used for PCR amplification in a 50μL reaction with 10μM primers 5’-AATGATACGGCGACCACCGAGATCTACACTCTTTCCCTACACGA and 5’-CAAGCAGAAGACGGCATACGAGAT and NEB Next high-fidelity 2X mix (NEB), using 98°C 30sec; 18 cycles of 98°C 10sec, 58°C 40sec, 72°C 30sec; 72°C 5min, followed by 0.9X AMPure purification. Entire eluate was then run on a 2% E-gel (ThermoFisher) and fragments 200pb-500bp were gel extracted. Library quality and quantity were estimated with Bioanalyzer and Qubit assays. Libraries were sequenced on a Next-seq 500 using 75bp single end.

### ChIP-Exo

For fixation, 10 million adherent mESCs were incubated in 2% formaldehyde-10%FBS in PBS for 10 min at room temperature, quenched by adding glycine to 0.125M, washed with 0.125M glycine in PBS, scraped, pelleted, snap frozen on dry ice and stored at −80°C.

Procedure was based on (Luna-Zurita et al., 2016) with modification. Chip procedure was the same as for Chip-seq except that no spike-in antibody was used and washes consisted in 6 iterations of RIPA buffer (HEPES pH7.6 50mM, EDTA 1mM, Sodium Deoxycholate 0.7%, NP40 1% and LiCl 0.5M) followed by two iterations of Tris-HCl pH8. End repair was immediately followed by resuspending the DNA-antibody-bead matrix with 1mM ATP, 100μM dNTPs, 15U T4 DNA polymerase (NEB), 5U Klenow large fragment DNA polymerase (NEB) and 50U T4 PNK (NEB) in 1X NEB buffer 2 and incubating at 30°C for 30min. After two RIPA and two Tris-Cl pH8 washes ligation of p7 adapters was performed by resuspending the beads in 100μL of 1mM ATP, 150pmol p7 adapter and 2000U T4 DNA ligase (NEB) in 1X NEB buffer 2 and incubating at 25°C for 60min. p7 adapters were prepared by mixing the following HPLC purified oligos at 10μM final in 10M Tris-Hcl pH8, 50m NaCl, 1M EDTA: 5’Phos-GTGACTGGAGTTCAGACGTGTGCTCTTCCGATC 3’ and 5’-GATCGGAAGAGCACACGTCT. After two RIPA and two Tris-Cl pH8 washes Nick repair was performed by resuspending the beads in 100μL of 150μM dNTPs, 15U Phi29 polymerase (NEB) in 1X Phi29 polymerase buffer (NEB) and incubating at 30°C for 20min. After two RIPA and two Tris-Cl pH8 washes lambda exonuclease digestion was performed by resuspending the beads in 100μL 1X Lambda exonuclease buffer supplemented with 10U lambda exonuclease (NEB) and incubating at 37°C for 30min. After two RIPA and two Tris-Cl pH8 washes RecJf exonuclease digestion was performed by resuspending the beads in 100μL 1X RecJf exonuclease buffer supplemented with 30U lambda exonuclease (NEB) and incubating at 37°C for 30min. After two RIPA and two Tris-Cl pH8 washes DNA was finally eluted by adding 100μL of 50mM NaHCO3, 1%SDS and incubating at 65°C for 30min. Supernatent was collected and supplemented with 1μL of 10mg/mL RNAse A, incubated at 37°C for 30min. 1μL of 20mg/mL Proteinase K and 12μL of 5M NaCl was then added and samples were reverse-crosslinked by incubation at 65°C overnight.

DNA was then purified using AMPure XP beads at a ratio 1.8X to sample and eluted in 20μL Tris-HCl pH8. DNA was then denatured by incubation at 95°C for 5min and immediate transfer on ice. Second strand was then synthesized by adding 5pmol of P7 primer (5’-GACTGGAGTTCAGACGTGTGCT) in 50μL total of 1X Phi29 buffer (NEB) and incubating at 65°C for 5 min then 30°C for 2min, followed by addition of 10U Phi29 polymerase and 1μL of 10M dNTPs and incubation at 30°C for 20min and 65°C for 10 minutes. Following AMPure XP purification (1.8X) and elution in 20μL ligation of p5 adapter was performed by incubation with 15pmol p5 adapter, 2000U T4 ligase in 1X T4 ligase buffer (NEB) in 50μL total at 25°C for 60min then 65°C for 10min. p5 adapters were prepared by mixing the following HPLC purified oligos at 10μM final in 10M Tris-Hcl pH8, 50m NaCl, 1M EDTA: 5’-AGATCGGAAGAGCG and 5’-TACACTCTTTCCCTACACGACGCTCTTCCGATCT. Following AMPure XP purification (1.8X) and elution in 20μL PCR amplification with indexed primers was performed using the NEB Next high-fidelity 2X PCR Master Mix with 25μM primers in 50μL and using 98°C for 30sec, 18 cycles of 98°C for 10sec, 65°C for 30sec and 72°C for 30sec, followed by 72°C for 5min. PCR primer sequences are 5’-AATGATACGGCGACCACCGAGATCTACACTCTTTCCCTACACG*A and 5’-CAAGCAGAAGACGGCATACGAGATNNNNNNGTGACTGGAGTTCAGACGTGTGC*T where * represents a phosphothiorate bond and NNNNNN is a Truseq index sequence. Following AMPure XP purification (0.9X) and elution in 20μL libraries were loaded on a 2% E-gel (ThermoFisher) and fragments 200pb-500bp were gel extracted. Library quality and quantity were estimated with Bioanalyzer and Qubit assays. Libraries were sequenced on a Hi-seq2000 or 4000 using 75bp single end.

### RNA-Seq

Total RNA was prepared by Ethanol precipitation as described in Jay & Ciaudo 2013. Six to ten million adherent mESCs where washed with PBS and lysed directly with Trizol (Thermofisher), transferred into a 15mL conical tube, vortexed, supplemented with 1.6mL Chloroform, vortexed again and centrifuged at 3200g at 4°c for 15 min. Upper phase was mixed with and equal volume of isopropanol and spun at 3200g 4°C 30 min. Pellet was washed with 70% ethanol, air dried and resuspended in 100μL water. 10μg total RNA was used with the DNAse turbo kit (Ambion) in 50μL with 1μL DNAse. To purify polyA+ species 10μg DNAse treated RNA was heated at 65°C 5 min, transferred on ice, mixed with 20μL oligodT(25) magnetic beads (ThermoFisher) prewashed and resuspended in 45μL binding buffer and incubated 15min at room temperature. After two 200μL washes beads were resuspended in 10μL Tris pH 7.5, heated at 75°C for 2 min and eluate was immediately subjected to a second round of purification using 10μL beads per sample and eluting in 20μL – resulting in 30-100ng RNA. RNA-seq library were constructed using the NEBNext ultra (non-directional) RNA library kit for Illumina using 10ng polyA+ RNA as input and 12-15 PCR cycles. Library concentrations were estimated using Bioanalyzer (Agilent) and Qubit (ThermoFisher) assays, pooled and sequenced on a Next-seq instrument (Illumina) using 1.8pM, 75bp paired-end.

### Chromosome Conformation Capture Carbon-Copy (5C)

We made substantial improvements over previously published protocols (Dostie et al. 2006), incorporating *in situ* (in nuclei) ligation (Rao et al. 2014), circumventing the need for phenolchloroform purification and adopting a single-PCR strategy to construct 5C-sequencing libraries from the 3C template. These changes enable proceeding through the 3C protocol in a single tube per sample, allow handling of over 20 samples in parallel, reduce the amount of cells needed by a factor 5 to 10 and cut down the time needed to complete the protocol from 8 days (Nora et al. 2012) to 4 days.

10 million adherent mESCs were fixed as described for Chip-exo except that 2% formaldehyde was used. For 3C, 5 million cells were Lysed in 1mL 10mM Tris-HCl pH8.0, 10mM Nacl 0.2% NP40 for 15 min, pelleted at 4°C and washed twice with 1mL ice cold 1X NEB buffer 2. Cells were then resuspended in a 1.5mL tube in 400L 0.1% SDS in 1X NEB buffer 2 at room temperature, incubated at 65°C for 10min, cooled, supplemented with 44μL 10% Triton X-100, incubated at 37°C for 15 min. 1000U of HindIII (high-concentration, NEB) was then added for overnight incubation in a thermomixer at 800rpm. Cells were then incubated at 65°C 20min, cooled at room temperature and supplemented with 800μL of 50μM Tris-HCl pH7.5, 10mM MgCl2, 10mM DTT, 1% Triton X-100, 0.1mg/mL BSA, 1mM ATP and 10U T4 ligase (ThermoFisher cat 15224017). After 4h incubation at 25°c in a thermomixer at 800rpm cells were centrifuged at 1000rpm, resuspended in 500μL of 1% SDS with 1μg Proteinase K in 1X TE buffer, incubated at 55°C for 30min, supplemented with 50μL of 5M NaCl and incubated at 65°C overnight. DNA was then purified by adding 500μL isopropanol and incubating at −80°c for 30min following by centrifugation at 18,000g at 4°C, one 70% Ethanol wash, air drying and resuspension in 50μL 1X TE buffer, followed by incubation with 10μg RNAse at 37°C.

For 5C-sequencing we used the set of oligonucleotides described in Nora et al. 2012 that we pooled omitting the ones that were previously found to produce aspecific ligation (**Supplementary table 5**). 3C template were quantified using gel electrophoresis or the PicoGreen assay (ThermoFisher). Two to four 20μL 5C annealing reactions were assembled in parallel, each using 1μg 3C template, 1μg Salmon Sperm (ThermoFisher), 10fmol of each 5C oligonucleotide in 1X NEB buffer4. For neural progenitor cells and astrocytes 4μg of 3C template was used per 20μL annealing reaction. Samples were denatured at 95°C for 5 minutes and incubated at 48°C for 12-16h. 20μL of 1X Taq ligase buffer with 5U Taq ligase were added to each annealing reaction followed by incubation at 48°C 1h and 65°C 10 min. Negative controls (no ligase, no template, no 5C oligonucleotide) were included during each experiments to ensure the absence of contamination.

To fuse Illumina-compatible sequences 5C libraries were directly PCR amplified with primers annealing to the universal T3/T7 portion of the 5C oligonucleotides (underlined) and harboring 5’ tails containing Illumina sequences *(Italic):*

5C-PCR_FOR:

5’*AATGATACGGCGACCACCGAGATCTACACTCTTTCCCTACACGACGCTCTTCCGATCT*<u>ATTAACCCTCACTAAAGGGA</u>

5C-PCR_REV:

5’*CAAGCAGAAGACGGCATACGAGATnnnnnnGTGACTGGAGTTCAGACGTGTGCTCTTCCGATCT*<u>TAATACGACTCACTATAGCC</u>

Where nnnnnn denotes a 6-bp Truseq index sequence (Illumina) for multiplexing.

For this each 5C ligation reaction was used to template two parallel PCRs (so 4-8 PCRs total), using per reaction 6μL of 5C ligation with 1.125U Amplitaq gold (ThermoFisher) in 1X PCR buffer II, 1.8mM MgCl2, 0.2 dNTPs, 1.25μM 5C-PCR_FOR and 5C-PCR_REV primers in 25μL total. Cycling conditions were 95°C 9 min, 25 cycles of 95°C 30sec, 60°C 30sec, 72°C 30 sec followed by 72°C 8min. PCR products from the same 3C sample were pooled and purified using the PCR purification MinElute kit (Qiagen) and run on a 2.5% agarose electrophoresis. 5C libraries (231bp) were then excised and purified with the Gel extraction MinElute kit (Qiagen). Library concentrations were estimated using Bioanalyzer (Agilent) and Qubit (ThermoFisher) assays, pooled and sequenced on a Next-seq instrument (Illumina) using 1.2 to 1.5pM and 20-40% PhiX, 92bp single end.

### Hi-C

Hi-C was performed as described (Lieberman-Aiden et al., 2009; Naumova et al., 2013). 25 million 2% formaldehyde cross-linked cells were incubated in 1000 μl of cold lysis buffer (10 mM Tris-HCl pH8.0, 10 mM NaCl, 0.2% (v/v) Igepal CA630, mixed with 10 μl protease inhibitors (Thermofisher 78438) immediately before use) on ice for 15 minutes. Next, cells were lysed with a Dounce homogenizer and pestle A (KIMBLE Kontes # 885303-0002) by moving the pestle slowly up and down 30 times, incubating on ice for one minute followed by 30 more strokes with the pestle. The suspension was centrifuged for 5 minutes at 2,000 g at RT using a table top centrifuge (Centrifuge 5810R, (Eppendorf). The supernatant was discarded and the pellet was washed twice with ice cold 500 μl 1x NEBuffer 2.1 (NEB). After the second wash, the pellet was resuspended in 1x NEBuffer 2 in a total volume of 250 μl and split into five 50 μl aliquots. Next, 312 μl 1x NEBuffer 2 was added to each aliquot. Chromatin was solubilized by addition of 38 μl 1% SDS per tube and the mixture was resuspended and incubated at 65°C for 10 minutes. Tubes were put on ice and 44 μl 10% Triton X-100 was added. Chromatin was subsequently digested by adding 400 Units HindIII (NEB) at 37°C for overnight digestion with alternating rocking. Digested chromatin solutions were spun shortly and transferred to ice. One tube was kept separate and used for generating a 3C control library as described (Naumova et al., 2013). The chromatin samples in the remaining four tubes were used for generating Hi-C libraries and were treated as follows: The HindIII DNA ends were filled in and marked with biotin by adding 60 μl fill-in mix [1.5 μl 10 mM dATP, 1.5 μl 10 mM dGTP, 1.5 μl 10 mM dTTP, 37.5 μl 0.4 mM biotin-14-dCTP (ThermoFisher #19518-018), 6 μl 10x NEBuffer 2.1, 2 μl water and 10 μl 5U/μl Klenow polymerase (NEB M0210L)] followed by incubation at 37°C for 80 minutes in a thermomixer. Klenow polymerase was inactivated by adding 96 μl 10% SDS followed by incubation at 65°C for 30 minutes. Tubes were then placed on ice immediately afterwards. The content of each of the tubes was transferred to 15 ml conical tube containing 7.58 ml ligation mix [820 μl 10% Triton X-100, 758 μl 10x ligation buffer (500 mM Tris-HCl pH7.5, 100 mM MgCl2, 100 mM DTT), 82 μl 10 mg/ml BSA, 82 μl 100 mM ATP and 5.84 ml water]. 50 μl 1U/μl T4 DNA ligase (Invitrogen #15224) was added and ligation was performed at 16°C for 4 hours. DNA was then purified as follows. 50 μl 10 mg/ml Proteinase K (ThermoFisher # 25530-031) was added to each tube and samples were incubated at 65°C for 4 hours followed by a second addition of 50 μl 10 mg/ml Proteinase K solution, followed by overnight incubation at 65°C. Tubes were cooled to RT and transferred to 50 ml conical tubes. The DNA was extracted by adding an equal volume of phenol pH8.0:chloroform (1:1) (Fisher BP1750I-400), vortexing for 3 minutes and spinning for 10 minutes at 4,000 rpm in a table top centrifuge (centrifuge 5810R, Eppendorf). The supernatants were transferred to new 50 ml conical tubes. Another extraction was performed with an equal volume of phenol pH8.0:chloroform (1:1). After vortexing and centrifugation for 10 minutes at 4,000 rpm, all four supernatants of the Hi-C samples were pooled into a single 250ml centrifuge tube and the volume was brought to 40 ml with 1x TE buffer (10 mM Tris pH8.0, 1 mM EDTA). To precipitate the DNA, 4 ml 3M Na-acetate pH5.0 was added, mixed well and then 100 ml of ice cold 100% ethanol was added. The volume of the 3C control sample was brought to 10 ml with TE. DNA precipitation was done by addition of 1 ml of 3M Na-acetate and 25 ml ice-cold 100% ethanol in a 35 mL centrifuge tube. Tubes were inverted slowly several times to mix the contents and then were incubated at least one hour at −80°C. Next, the tubes were spun at 4°C for 30 minutes at 16,000g AvantiTM J-25 Centrifuge (Beckman). The supernatants were discarded and DNA pellets were dissolved in 500 μl 1x TE buffer and transferred to a 0.5 mL AMICON^®^ Ultra Centrifugal Filter Unit – 0.5 ml 30K (UFC5030BK EMD Millipore) for desalting. Columns were spun at 14,000g for 10min, in a microfuge. The flow throughs were discarded. Columns were washed three times with 450 μl TE. After the final wash, the 3C library was dissolved in 25 μl TE; the Hi-C library was dissolved in 100 μl TE. Any RNA was degraded by incubation with 1 μl of 10 mg/ml RNAse A at 37°C for 30 minutes. The quality and quantity of 3C and Hi-C libraries were checked by running aliquots on a 0.8% agarose gel along with a 1 kb ladder (NEB #N3232S). Libraries should run as a rather discrete band with a molecular weight that is larger than 10 kb. With a successful biotin fill-in and marking of DNA ends, HindIII (AAGCTT) restriction sites get converted into NheI sites (GCTAGC). To test the efficiency of this process we used PCR to amplify a ligation product formed by two nearby restriction fragments followed by digestion with HindIII, NheI and by a double digestion with HindIII+NheI restriction enzymes. The relative efficiency of Hi-C ligation product formation and biotin fill-in was defined as the proportion of ligation product digested with NheI and varied from 50 to 80% in different Hi-C libraries. The following two pairs of primers were used: mGAPDH_1 and mGAPDH2.

mGAPDH_1 ATGGAGACCTGCCGCCGGCTCATCA

mGAPDH_2 CGTGCTGTGACTTCGCACTTTTCTGA

Next, Hi-C libraries were treated with T4 DNA polymerase to remove biotinylated ends that did not ligate (dangling ends). Eight reactions were assembled as follows: 5 μg of Hi-C library, 5 μl 10x NEBuffer 2.1, 0.5 μl 2.5 mM dATP, 0.5 μl 2.5 mM dGTP and 5 Units T4 DNA polymerase (NEB # M0203L) in a total volume of 50 μl Reactions were incubated at 20°C for 4 hours. The reaction was stopped by incubating 20min at 75°C. To desalt and concentrate the DNA, the reactions were pooled together and added on 0.5 mL AMICON^®^ Ultra Centrifugal Filter Unit – 0.5 ml 30K (UFC5030BK EMD Millipore). Columns were spun at 14,000g for 10min, in a microfuge. The flow through was discarded. Columns were washed twice with 450 μl TE. After the final wash, the Hi-C libraries was dissolved in 120 μl TE. The DNA was sheared to a size of 100-400 bp (with the majority of molecules around 200 bp) using a Covaris S2 instrument (Covaris, Woburn, MA). The settings were as follows: Duty cycle 10%, Intensity 5, Cycles per burst 200, Set mode – Frequency sweeping, Process time 60 sec per process, Cycles number 3. To enrich for DNA fragments of 100-300 bp an Ampure XP fractionation was performed (Beckman Coulter, A63881) and the DNA was eluted with 50 μL of water. The size range of the DNA fragments after fractionation was determined by running an aliquot on an agarose gel. The sheared DNA ends were repaired by addition of 7 μl 10x ligation buffer (NEB # B0202S), 7 μl 2.5 mM dNTP mix, 2.5 μl T4 DNA polymerase (NEB # M0203L), 2.5 μl T4 polynucleotide kinase (NEB #M0201S), 0.5 μl Klenow DNA polymerase (NEB #M0210S) and 5.5 μl water to the 45uL of DNA. The DNA was purified using Ampure beads (Beckman Coulter, A63881) and eluted in 32uL of TLE (10 mM Tris pH8.0, 0.1 mM EDTA (TLE buffer). Next, an ‘A’ was added to the 3’ ends of the end-repaired DNA by addition of 5 μl 10x NEBuffer2, 10 μl 1 mM dATP, 3 μl Klenow (exo-) (NEB #M0212L) and 16 μl water. The reaction was incubated at 37°C for 30 min followed by incubation at 65°C for 20 minutes to inactivate Klenow polymerase. The reactions were cooled on ice. All subsequent steps were performed in DNA LoBind tubes (Eppendorf #22431021) and each step was performed in a fresh tube. 50 μl of streptavidin Dynabeads (MyOne Streptavin C1 Beads, ThermoFisher #650-01) were washed twice with 400 μl Tween Wash Buffer (TWB) (5 mM Tris-HCl pH8.0, 0.5 mM EDTA, 1 M NaCl, 0.05% Tween20) by incubating for 3 minutes at RT with rotation, reclaiming against a magnetic separation rack for 1 minute and removing all supernatant. Next, reclaimed beads were resuspended in 400 μl 2x Binding Buffer (BB) (10 mM Tris-HCl pH8.0, 1 mM EDTA, 2 M NaCl) and combined with 400 μl Hi-C DNA from the previous step. The mixture was incubated at RT for 15 minutes with rotation. The supernatant was removed and the DNA-bound Streptavidin beads were washed once with 400 μl 1x BB. The beads were then washed with 100 μl 1x ligation buffer (Invitrogen 5x buffer), and then resuspended in 19 μl of 1x ligation buffer (NEB quick ligase, M2200S). Ligation reaction was set-up as follows: 19 μl Hi-C library on beads, 6 μl Illumina paired end adapters (Illumina), 10 μl 2x quick ligation buffer (NEB), 1 μl quick DNA ligase (NEB quick ligase, M2200S). The reaction was incubated at RT for 15min. The beads with bound ligated Hi-C DNA were collected by holding against a magnetic separation rack and were then washed twice with 400 μl 1x TWB, once with 200 μl 1xBB and once with 200 μl 1x NEBuffer2 to remove non-ligated Paired End adapters. The beads were resuspended in 20 μl 1x NEBuffer 2. Next, test PCR reactions were performed to determine the optimal number of PCR cycles needed to generate enough Hi-C library for sequencing. Four trial PCR reactions were set up, each containing 0.9 μl Dynabead-bound Hi-C library, Illumina PE1.0 and PE2.0 PCR primers (0.21 μl of each; 25 μM), 0.12 μl 25mM dNTPs, 0.3 μl Pfu Ultra II Fusion DNA polymerase (Agilent #600670), 1.5 μl 10x Pfu Ultra buffer and 11.76 μl water. The temperature profile during the PCR amplification was 30 seconds at 98°C followed by 5, 7, 9 or 11 cycles of 10 seconds at 98°C, 45 seconds at 65°C, 30 seconds at 72°C and a final 7-minute extension at 72°C. The PCR reactions were run on a 2% agarose gel and the minimal cycle number was determined that yielded sufficient DNA for sequencing. Typically, 6 cycles were chosen for amplification of Hi-C libraries. PCR was then performed in nine reactions with the remaining Dynabead-bound Hi-C library. The PCR product was run on a 2% agarose gel and smear 200-400bp to assess the DNA concentration. A final quality control was performed by NheI digestion of an aliquot of the final Hi-C library. Without NheI digestion, the DNA sizes of the libraries ranged from 300-400bp. After NheI digestion, the DNA sizes of the libraries shifted and ranged from 100-350bp. It indicated that the majority of the ligation products have been digested by NheI and validated that the libraries were mainly constituted of true ligation products. The libraries were sequenced using 50 bp paired end reads with a HiSeq2000 machine and HiSeq4000.

## QUANTIFICATION AND STATISTICAL ANALYSIS

### ChIP-seq Analysis

Fastq files were trimmed using the fastq-mcf program, aligned to the mm9 reference genome with bowtie2 (Langmead and Salzberg, 2012). Reads with a mapq score of 30 or greater were retained, using Samtools. Data used to generate the heatmaps presented in the manuscript were obtained by downsampling the number of reads to match the most shallow sample (for CTCF and H3K27me3 separately) and pooling the reads of each biological replicates. Heatmap visualization and integration with RNA-seq was performed using Easeq version 1.03 (Lerdrup et al., 2016). The Euler diagram was drawn using eulerAPE (Micallef and Rodgers, 2014). Chip-seq peaks were called on each replicate individually using all available reads. For peak calling we followed the guidelines described in (Thomas et al., 2016). For CTCF, which display focal enrichment, we used the Genome-wide Event finding and Motif discovery (GEM) method (Guo et al., 2012). For H3K27me3, which marks broad domains, we used the Baysian Change Point (BCP) method (Xing et al., 2012). The consensus peak list was obtained by retaining peaks that overlapped for at least 1bp between biological replicates. For exemple loci in figures S2 and S7 read depth-normalized tag densities were generated directly by the Easeq software using the “filled track” tool. The normalized tag density bigwig tracks used for visualization with the UCSC genome browser were generated by dividing into 20bp bins and a normalized tag density was calculated for each bin as follow:

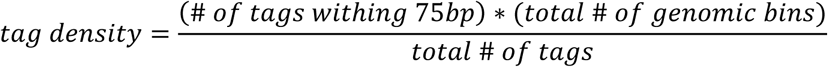

### ChIP-Exo Analysis

Analysis and footprint identification was carried out as described in (Luna-Zurita et al., 2016). The 5’-most position of reads that mapped to the reference strand and the 3’-most position of reads that mapped to the non-reference strand were identified for each read as the actual edges of each exonuclease-treated fragment. To identify broad regions of binding, bins with tag densities of greater than 100 were merged to generate a peak list for each sample. Within 1kb of each region, strand-specific single-base-resolution tag densities were calculated for each dataset by dividing each region into 1bp bins, then counting the number of tags within 5bp of each bin. For each region of binding, the footprint for each bound region was defined as the span from the peak position of ‘+’ strand binding to the peak position of ‘−’ strand binding as seen from the high-resolution tag densities.

### RNA-seq Analysis

Alignment and differential expression was performed on the BaseSpace environment version 1.0.0 (Illumina). Alignment was produced using STAR version 2.5.0a (Dobin et al., 2013) with default parameters except that novel transcript assembly was not performed. mm9 RefSeq was used as reference gene set and adapters were trimmed, Cufflinks version 2.2.1 (Trapnell et al., 2010)was used with fragment bias and multi-read correction with Bedtools version 2.17.0 (Quinlan and Hall, 2010). Differential expression analysis was analyzed using Cuffdiff (Trapnell et al., 2013) with default parameters within BaseSpace. Genes with an FPKM below 1.1 in all conditions were not considered in the differential expression analysis. Heatmap visualization and integration with Chip-seq and Chip-exo was performed using Easeq version 1.03 (Lerdrup et al., 2016). For integration with enhancer positions we took the enhancer list assembled by (Chen et al., 2012) with the same probability threshold (0.8). The super-enhancer list was retrieved from (Hnisz et al., 2013). FPKM provided in **Supplementary Table 3** are means from 3 independent biological replicates.

### CTCF Motif orientation analysis

First, we established a consensus list of CTCF ChIP-seq peak from the CTCF-AID line by retaining only the peaks identified in both replicates (overlap of at least 1bp between replicates). We then retrieved the DNA sequence from each peak using the TableBrowser tool of the UCSC genome browser, using the mm9 assembly. Each of these sequences were then searched for CTCF motifs using FIMO (Grant et al., 2011) with the CTCF position frequency matrix obtained from the JASPAR database, motif MA0139.1 and default parameters. Promoters of affected genes were not specifically enriched for tandem CTCF sites compared to their occurrence in CTCF ChIP-seq peaks genome-wide (around 1/3 of peaks have multiple CTCF motifs (Pugacheva et al., 2015)).

### Hi-C analysis

#### Mapping, filtering, and normalization of Hi-C data

We mapped the sequence of Hi-C molecules to reference mouse genome assembly mm9 using Bowtie 2.2.8 and the iterative mapping strategy, as described in (Imakaev et al., 2012; Lajoie et al., 2015). Upon filtering PCR duplicates and reads mapped to multiple or zero locations, we aggregated the reads pairs into 20kb and 100kb genomic bins to produce Hi-C contact matrices. For downstream analyses, data from biological replicates were pooled. Low-coverage bins were then excluded from further analysis using the MAD-max (maximum allowed median absolute deviation) filter on genomic coverage, set to three median absolute deviations. To remove the short-range Hi-C artifacts - unligated and self-ligated Hi-C molecules - we ignored the contacts mapping to the same or adjacent genomic bins in all downstream analyses. The filtered 20kb and 100kb contacts matrices were then normalized using the iterative correction procedure (IC), such that the genome-wide sum of contact probability for each row/column equals 1.0. Observed/expected contact maps were obtained by dividing each diagonal of a contact map by its chromosome-wide average value over non-filtered genomic bins. The compartment structure of Hi-C maps was detected using a modified procedure from (Imakaev et al., 2012). Compartments were quantified as the dominant eigenvector of the observed/expected 20kb and 100kb cis contacts maps upon subtraction of 1.0, as implemented in hiclib. Segmentation of eigenvectors into regions corresponding to active (A) and inactive (B) compartments was performed using a 2-state HMM model. The code for mapping, filtering and normalization analysis of Hi-C data is available at https://github.com/dekkerlab/cworld-dekker (lab of Job Dekker) and https://bitbucket.org/mirnylab/hiclib (lab of Leonid Mirny).

#### Insulation scores from Hi-C data

To local contact insulation analysis was based on the algorithm described in (Crane et al., 2015). For every 20kb bin, the insulation score was calculated as the total number of normalized and filtered contacts formed across that bin by pairs of loci located on the either side, up to 100kb away. The score was normalized by its genome-wide median. To find insulating boundaries, we detected peaks in log2-transformed insulation score track using the peakdet algorithm [Billauer E (2012). peakdet: Peak detection using MATLAB, http://billauer.co.il/peakdet.html]. Briefly, this algorithm seeks a sequence of local maxima and minima whose values differ by more than a pre-specified threshold (i.e. peak prominence). The detected minima in the insulation score correspond to a local depletion of contacts across the genomic bin, are then called as insulating boundaries. To find the optimal threshold for peak calling, we varied the peak calling threshold, and for each value compared the called boundaries with the loop anchoring regions detected in (Mumbach et al., 2016). This comparison revealed that at high threshold values, corresponding to stricter boundary selection, up to 62% of detected boundaries co-aligned with the previously detected loop anchors within +-1 20kb bin precision. As we lowered the boundary detection threshold, fewer added boundaries co-aligned with the loops; this analysis suggested the optimal threshold of 0.3, where the efficiency of loop anchor recall dropped 2-fold. Finally, we selected only the boundaries that had zero or one filtered 20kb bin in a 100kb range, since the presence of filtered bins affects insulation. We then used the same approach to call boundaries in all Hi-C samples. The boundaries detected in the auxin-treated sample within +-1 20kb bin from a position of a boundary in the untreated sample were called “residual” boundaries. To correlate the presence of boundaries with presence of active promoters or compartment transition we found all boundaries that had a PolII peak or a transition between A and B HMM compartment assignment, correspondingly; to account for the inaccuracy in boundary calls we allowed +/-1 20kb bin mismatch.

### Chromosome Conformation Capture Carbon-Copy (5C) analysis

#### Mapping and insulation scores from 5C matrices

Adapters were trimmed and aligment was performed using Bowtie2 against a pseudo-genome composed of all possible Forward-Reverse pairs of 5C oligonucleotides (Nora et al. 2012). Results were then transformed into a matrix table. Primers giving artefactual signal were removed using the code deposited by the lab of Job Dekker on Github https://github.com/dekkerlab/cworld-dekker. Heatmaps were generated using the my5C tools (Lajoie et al., 2009). For the insulation score analysis, the primer-based heatmaps were aggregated at 20kb resolution by calculating the median interaction frequency between primers belonging to all pairs of 20kb genomic bins. The aggregated maps were then filtered by removing the contacts in the first two diagonals and normalized using IC. The insulation score and boundary detection was then performed using the same method and parameters as described above for Hi-C maps.

#### Display of restriction fragment level 5C heatmaps

5C primers matrices were filtered as previously described methods (Sanyal, Dekker 2012). We detected and flagged all outlier (anchor) row/cols that are defined as having a having an aggregate (row/col) signal greater than or less than 1.5 * IQR (of the distribution of all row/col signals). We then took the union of all flagged (anchor) row/col outliers across all the 5C matrices, and removed these (anchor) row/cols from all datasets. 68 anchors were removed. Then, the matrices were balanced according to the ICE method developed for Hi-C (Imakaev, Mirny 2012. 5C cannot interrogate contacts between two restriction fragments harboring a forward oligonucleotide or a reverse oligonucleotide. In order to display intelligible heatmaps we interpolated the uninterrogated forward-forward and reverse-reverse pixels by the median of the eight pixels surrounding it, producing smoothed matrices using the my5C tools (Lajoie et al., 2009).

### 3D-DNA FISH analysis

3D distance measurement was performed on ImageJ using the scripts described in (Nora et al., 2012).

### Data and software accessibility

Software from this study has been previously published as detailed under “QUANTIFICATION AND STATISTICAL ANALYSIS.”

### Data Resources

Raw and processed sequencing data reported in this paper have been submitted to GEO, accession number pending.

Plasmids can be requested through Addgene.

## SUPPLEMENTARY FIGURES

**Figure S1.**
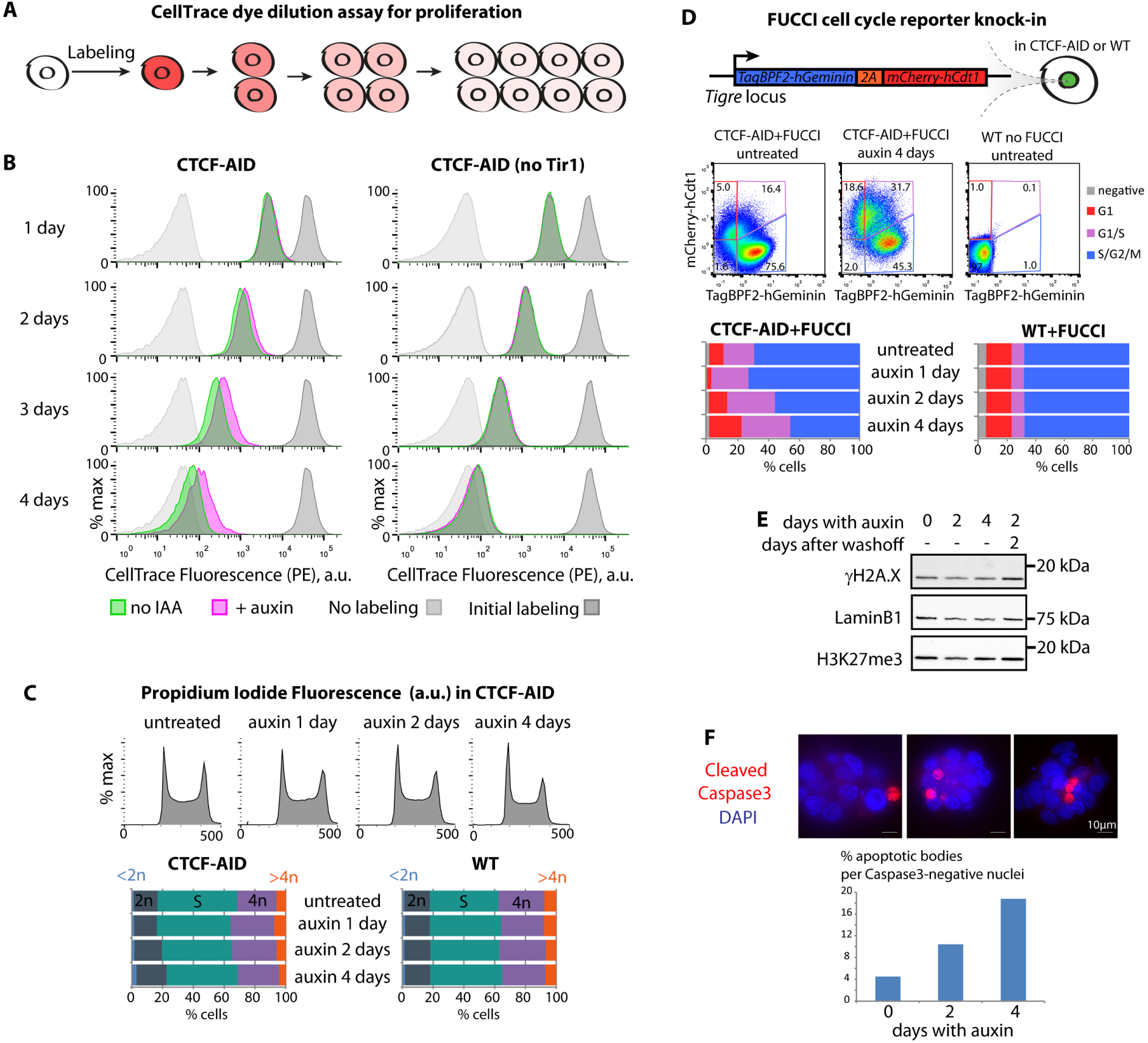
Characterization of the CTCF-AID mESCs. (A) Principle of the CellTrace dye dilution assay for proliferation (B) Flow cytometry of dilution kinetics of the CellTrace dye indicates that auxin-treated CTCF-AID mESCs keep proliferating after two days of CTCF depletion and slow down afterwards. Auxin does not trigger any proliferation defect in CTCF-AID mESCs lacking the Tir1 F-box protein transgene. (C) Propidium iodide staining indicates CTCF depleted mESCs are not blocked in any specific stage of the cell cycle and do not become aneuploidy. (D) A tagBFP2/mCherry FUCCI cassette was created and knocked in in CTCF-AID or WT mESCs. Auxin treatment only leads to a slight increase of the G1 FUCCI-signal after 4 days of CTCF depletion, confirming that loss of CTCF does not block cell cycle progression overall. (E) CTCF depleted did not show increased DNA damage as monitored by western blot, and displayed overall constant bulk H3K27me3 levels. LaminB1 used as loading control (F) CTCF depletion did not lead to massive apoptosis, although number of dying cells increase after long depletion.

**Figure S2.**
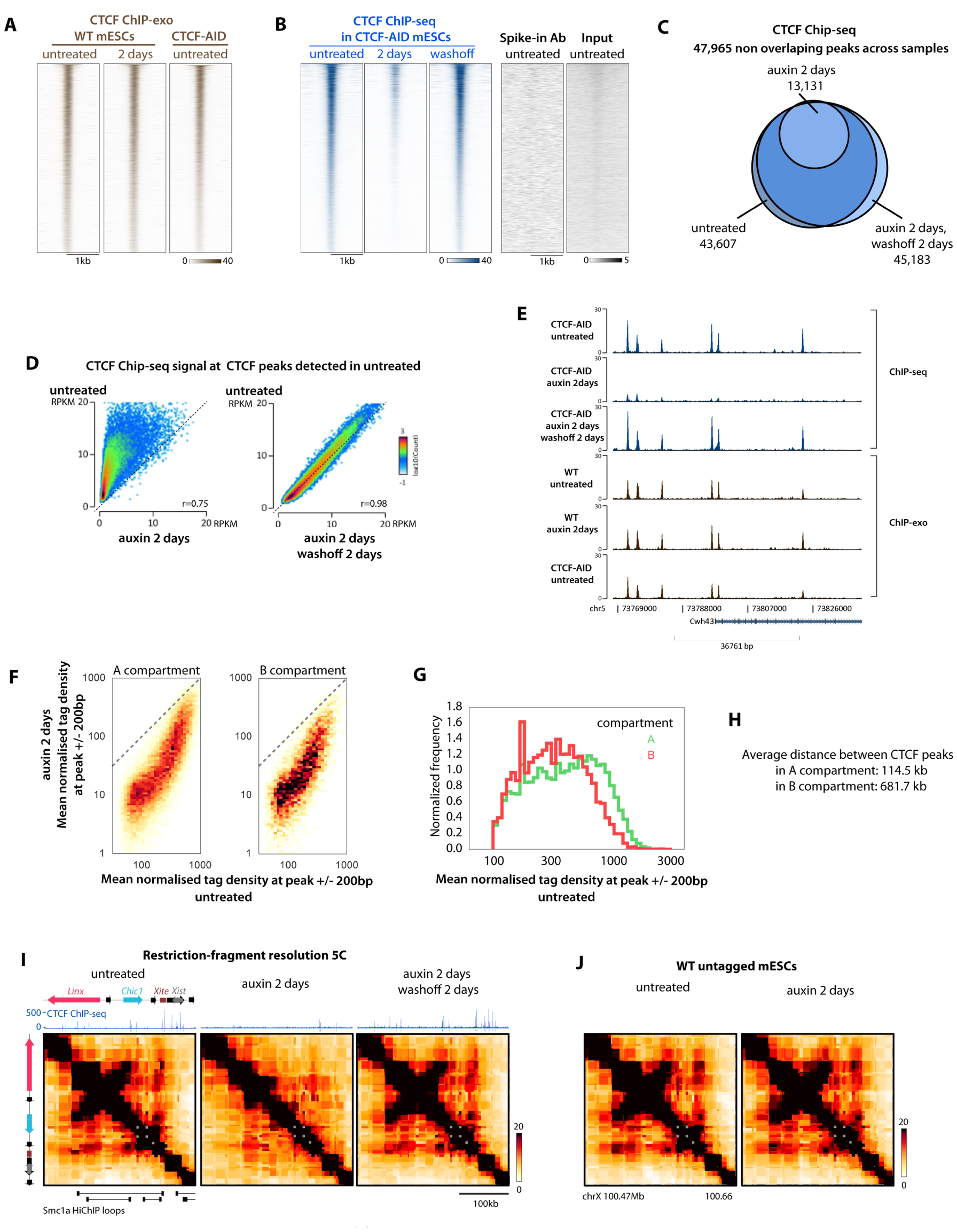
CTCF-ChIP seq analysis and Chromosome Conformation Capture Carbon-Copy (5C) (A) 5C at the *Xic* confirms that chromatin loops do not accumulate at CTCF peaks after CTCF depletion and are reaquired upon CTCF resoration. (B) Auxin treatment of WT cells has no effect on chromatin folding (C) CTCF ChIP-exo signal at CTCF ChIP-seq peaks detected in untreated CTCF-AID cells. Auxin treatment of WT cells has no effect on CTCF binding. Tagging with the CTCF-AID-eGFP does not disrupt CTCF binding pattern. (D) Auxin treatment of CTCF-AID cells dramatically reduces CTCF enrichment at peaks detected in untreated cells and is fully reversible after washoff (E) Easeq Genome browser visualization of an example locus. A subset of CTCF ChIP-seq peaks are still detected, but of low intensity, after depletion and are restored in strength after washoff (F) Loss of ChIP-seq signal upon CTCF depletion is equivalent in the A and B genomic compartment as defined by Hi-C. (G) A compartment tends to have stronger CTCF ChIP-seq peaks than B compartment (H) CTCF binding is 5-fold denser in the A compartment than in B. (I) Restriction-fragment level interpolated visualization of 5C around the *Linx-Chic1-Xite* loops. CTCF depletion disrupts CTCF binding and underlying loops while CTCF recovery re-stablishes binding and chromatin contacts. (J) Auxin treatment in itself does not perturb the accumulation of chromatin loops in WT untagged mESCs.

**Figure S3.**
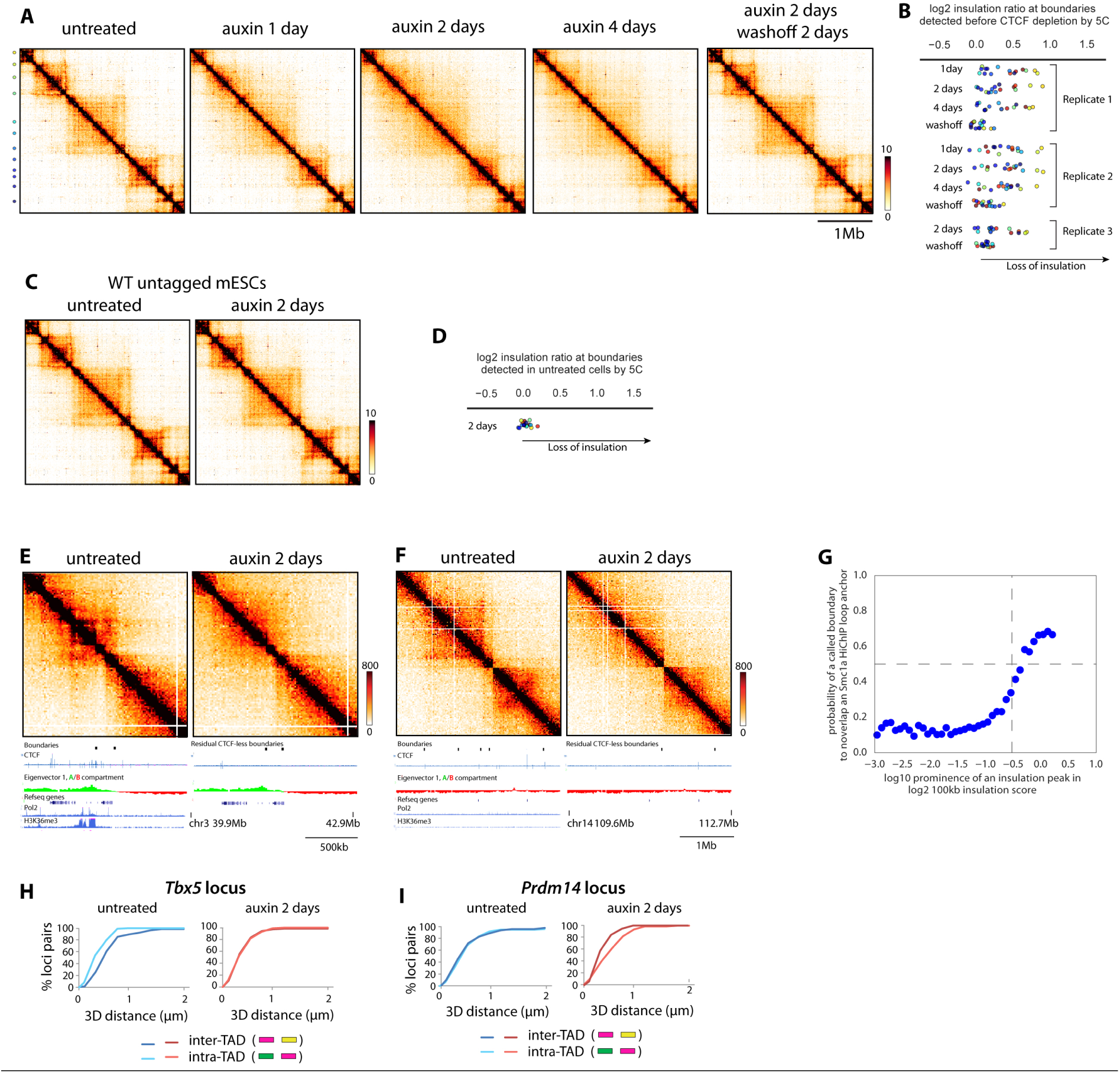
Supporting data regarding loss of TAD insulation upon CTCF depletion. (A) Restriction-fragment level interpolated visualization of 5C at the *Xic.* Color dots denote TAD boundaries. (B) Insulation score ratio between treated and untreated cells at boundaries detected in untreated cells, highlighting that a subset of TAD boundaries rapidly loose insulation upon CTCF depletion (C-D) Auxin treatment has no effect on TAD insulation in WT untagged cells (C) Hi-C snapshot illustrating that a subset of boundaries resist CTCF depletion Shown is an example region harboring boundaries that resist CTCF depletion. The left one is associated with a strong promoter and the other one with a A/B compartment transition. (D) Hi-C snapshot illustrating that a small subset of boundaries retain strong insulation after depletion without being associated with transcription or compartment transition (E) Probability of calling a TAD boundary at Smc1a HiChIP loop as a function of the local prominence of the insulation score calculated at 100kb with our Hi-C. We chose the threshold (0.3) below which improvement in retrieving Sm1a HiChIP loop is below 50% (see **methods**). (F) Replot of the DNA FISH data presented in figure 3D illustrating that after CTCF depletion inter-TAD 3D distances becomes equivalent to intra-TAD when probe pairs are equally spaced on the chromosome and not overlapping boundaries. (G) Replot of the DNA FISH data presented in figure 3D illustrating that probe pairs partially overlapping boundaries (green and yellow) become more separated after CTCF depletion

**Figure S4.**
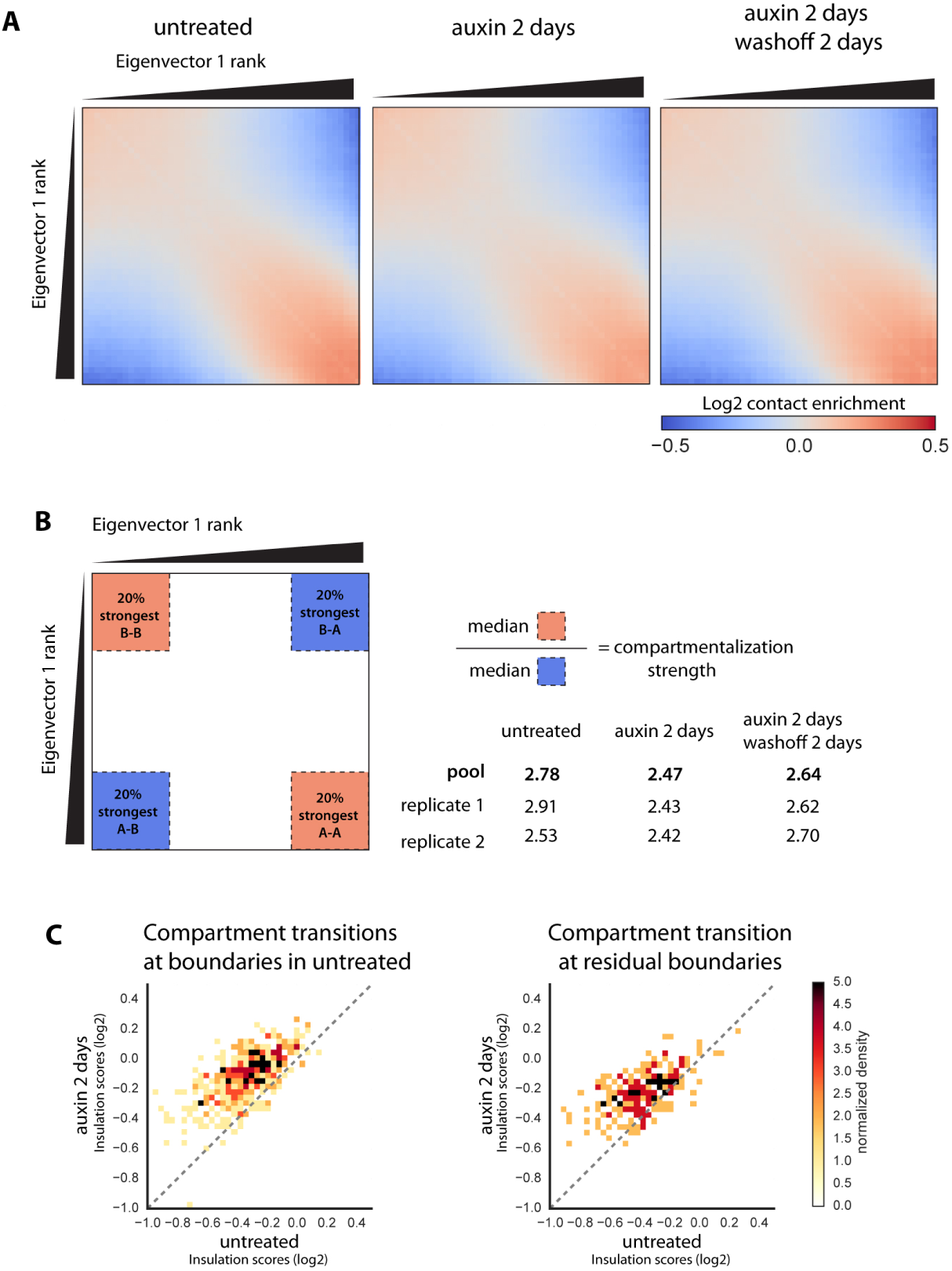
Large-scale chromosome folding is largely unaffected by CTCF depletion. (A) *cis*-Eigenvector 1 values in 100kb genomic bins are ranked and pairwise enrichment of Hi-C contacts between each of the 50 ranks are calculated (pooled replicates). Genomic regions with similar ranks of Eigenvector 1 values display more Hi-C contact while regions of opposite ranks are depleted (see methods). This trend is conserved overall after CTCF depletion or restoration. (B) Compartmentalization strength is only mildly affected by CTCF depletion (C) Scatter plot of the insulation score of genomic elements that are transitions between A/B compartments and TAD boundaries before CTCF depletion (left) or after (right), highlighting that insulation is also weaker at these compartments transition after loss of CTCF. Note that strong boundaries have the lowest insulation scores.

**Figure S5.**
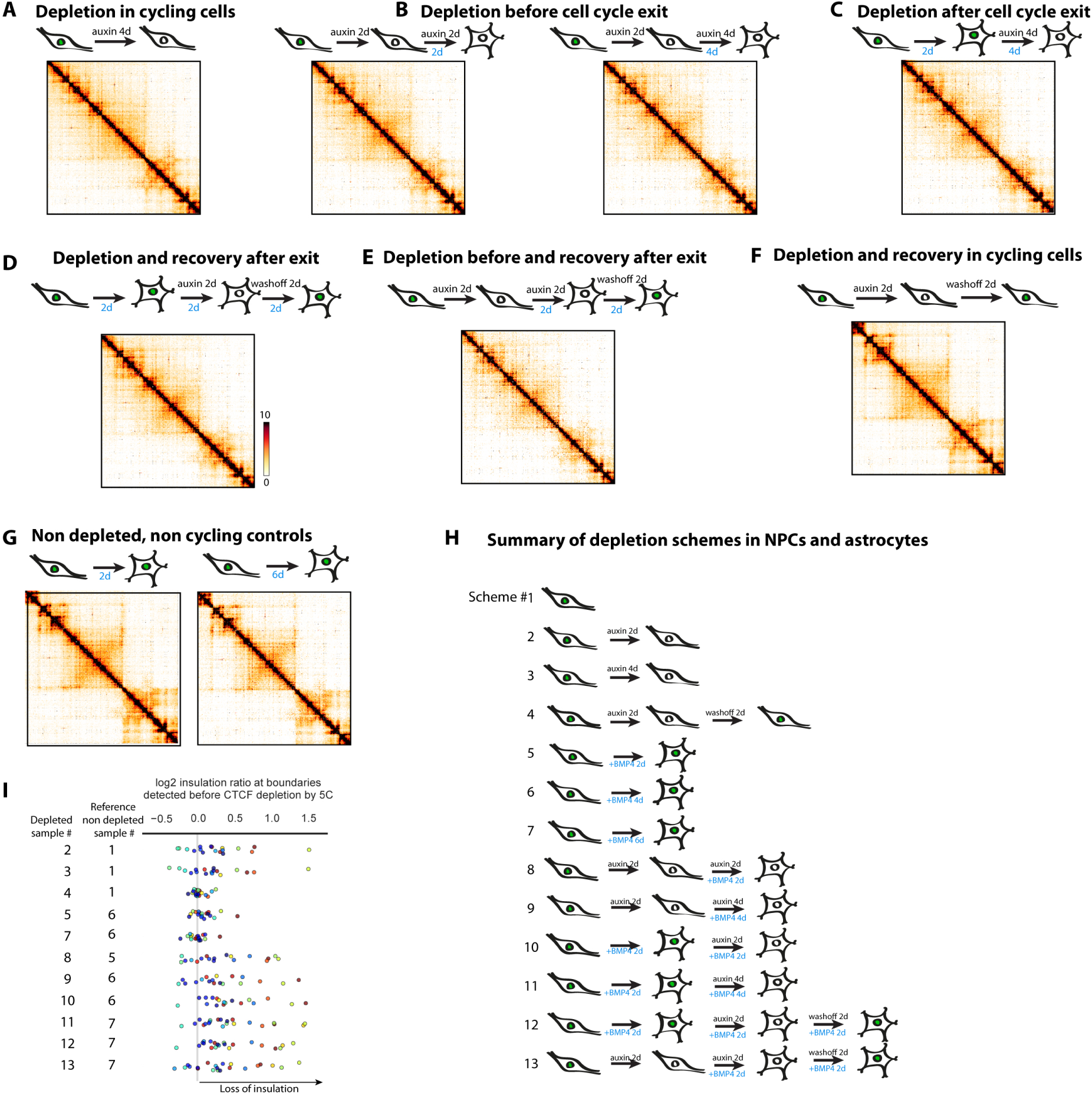
CTCF is required for proper TAD folding in differentiated and non-cycling cells. (A-G) Restriction-fragment level 5C interpolated heatmaps highlighting that auxin treatment of CTCF-AID NPCs and astrocytes disrupts TAD insulation, irrespective of the timing of CTCF depletion to cell-cycle exit. Blue = time after adding BMP4 to convert NPCs into astrocytes. (H) Summary of all experiments with NPCs and astrocytes. (I) Quantification of insulation loss from the 5C in all NPC and astrocyte samples.

**Figure S6.**
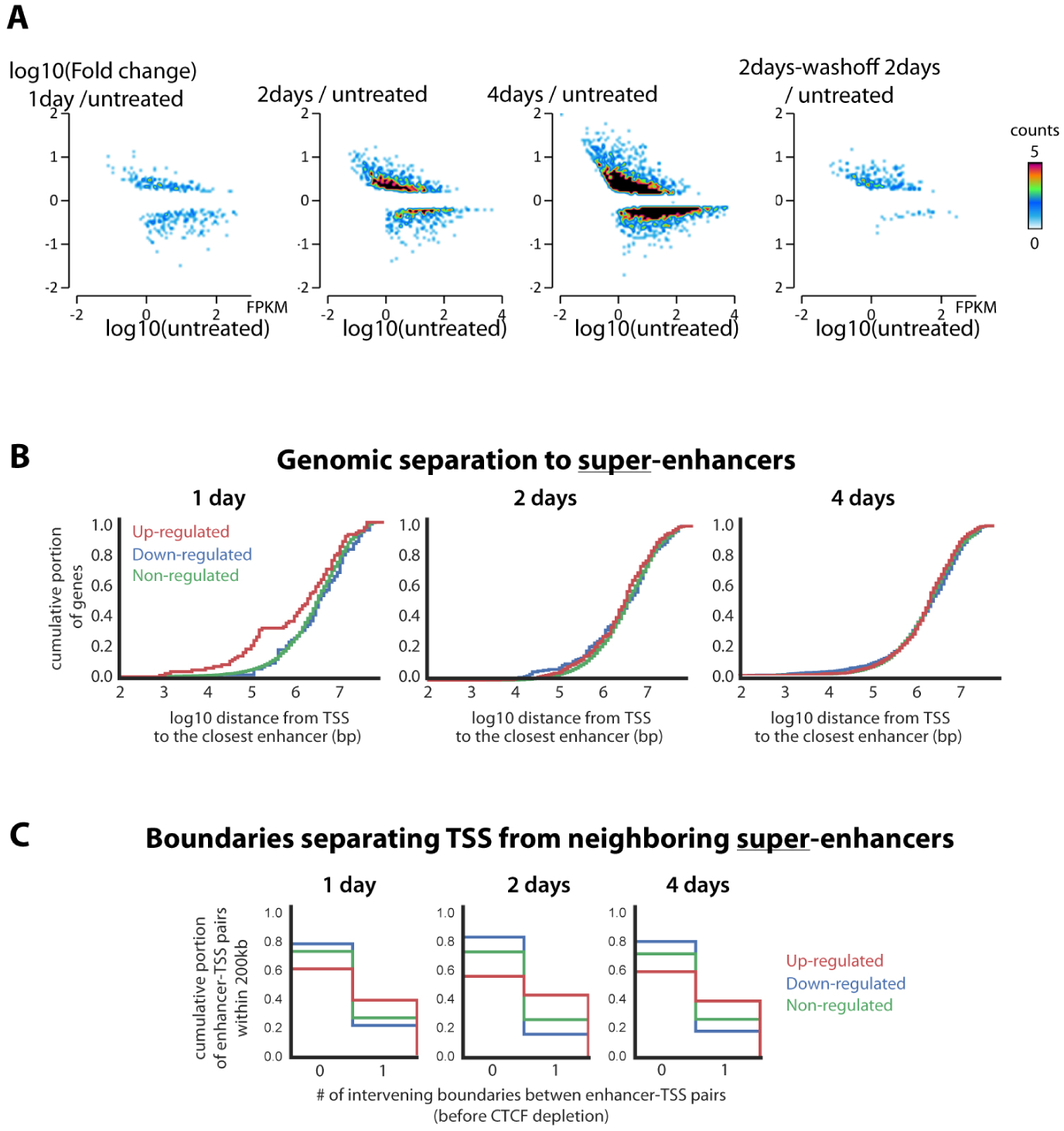
Supporting analyses of the RNA-seq after CTCF depletion. (A) Scatter plot of the fold change in treated versus untreated cells as a function of the expression level in untreated cells. Up and down-regulation are observed for genes with a wide-range of initial expression levels. Misregulation is not restricted to lowly-expressed genes. (B-C) Same analysis as in 6D-E but focused on super-enhancers active in mESCs.

**Figure S7.**
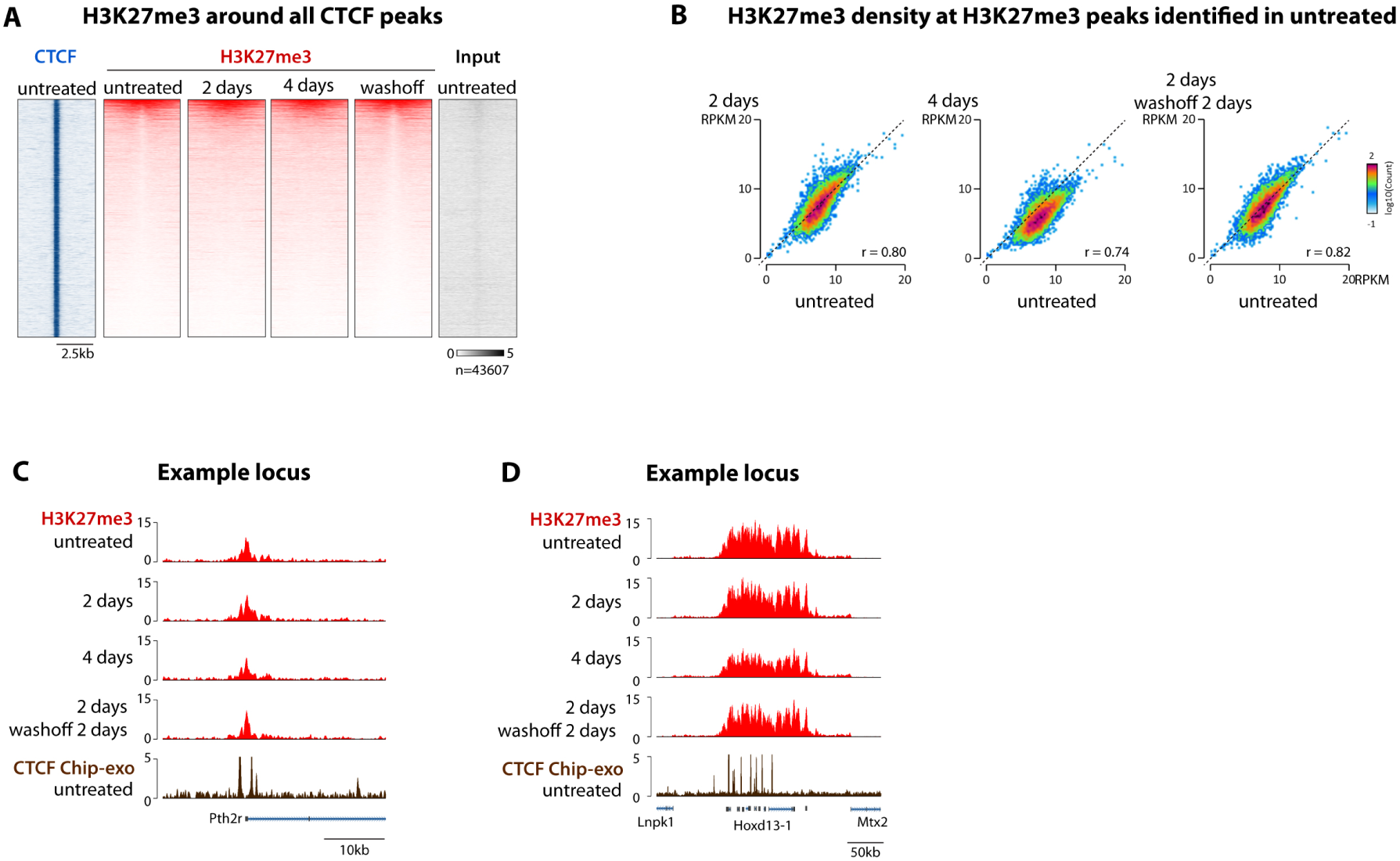
Additional analyses of H3K27me3 patterns after CTCF depletion. (A) CTCF and H3K27me3 ChIP-seq centered at all CTCF peaks detected in untreated cells. A small subset is embedded in large H3K27me3 regions with a dip at the CTCF site. This dip disappears upon CTCF depletion and reappears after CTCF restoration. This suggests that nucleosomes become able to cover the previously occupied CTCF site when CTCF binding is lost. (B) Overall H3K27me3 levels at H3K27me3 ChIP-seq peaks are unaffected after two day, become slightly lower after 4 days of depletion and are readjusted upon CTCF restoration. (C-D) Easeq Genome browser visualization of an example locus illustrating H3K27me3 does not spread beyond flanking CTCF sites upon CTCF depletion in mESCs.

## SUPPLEMENTARY TABLES

**Supplementary Table 1**. sgRNA sequences

**Supplementary Table 2**. List of sequencing experiments and mapping summary statistics

**Supplementary Table 3**. Mean RNA-seq FPKM from three biological replicates

**Supplementary Table 4**. ChIP-seq peaks that overlap between two biological replicates

**Supplementary Table 5**. List and sequence of 5C oligonucleotides

**Supplementary Table 6**. List of 20kb bins with insulation scores, boundary calls and compartment assignment

**Supplementary Table 7**. CTCF site orientation in ChIP-seq peaks

